# Hypothalamic CRH Neurons Modulate Sevoflurane Anesthesia and The Post-anesthesia Stress Responses

**DOI:** 10.1101/2023.08.02.551607

**Authors:** Shan Jiang, Lu Chen, Wei-Min Qu, Zhi-Li Huang, Chang-Rui Chen

**Author notes:** Corresponding author. (C-R Chen); (Z-L Huang). These authors contributed equally to this work (Shan Jiang, Lu Chen).

## Abstract

General anesthesia is an indispensable procedure necessary for safely and compassionately administering a significant number of surgical procedures and invasive diagnostic tests. However, the undesired stress response associated with general anesthesia (GA) causes delayed recovery and even increased morbidity in the clinic. Here, a core hypothalamic ensemble, corticotropin-releasing hormone neurons in the paraventricular nucleus of the hypothalamus (PVH^CRH^ neurons), is discovered to play a role in regulating sevoflurane GA. Chemogenetic activation of these neurons delay the induction of and accelerated emergence from sevoflurane GA, whereas chemogenetic inhibition of PVH^CRH^ neurons accelerates induction and delays awakening. Moreover, optogenetic stimulation of PVH^CRH^ neurons induce rapid cortical activation during both the steady and deep sevoflurane GA state with burst-suppression oscillations. Interestingly, chemogenetic inhibition of PVH^CRH^ neurons relieve the sevoflurane GA-elicited stress response (e.g., excessive self-grooming and elevated corticosterone level). These findings identify PVH^CRH^ neurons modulate states of anesthesia in sevoflurane GA, being a part of anesthesia regulatory network of sevoflurane.

**Impact statement:** Discovery of critical brain nodes regulating sevoflurane general anesthesia and the post-anesthesia stress responses.

## 1. Introduction

In the past century, volatile general anesthetics have gained worldwide popularity in clinics owing to their favorable properties, including a more pleasant odor, higher potency, low toxicity, and rapid recovery ^1^. General anesthesia is a combination of behavioral and physiological states induced and maintained primarily by pharmacologic agents. As an ideal volatile anesthetic agent, sevoflurane is commonly used for rapid and smooth induction and maintenance for general anesthesia (GA) ^2^. However, undesirable excessive stress responses of patients toward sevoflurane GA may result in prolonged rehabilitation and long-term prognosis ^3^.

There is increasing evidence supporting the ‘shared circuits hypothesis’ of GA and sleep, in which different GA drugs may exert hypnotic effect through a shared brain network with wake-sleep regulation ^4^. The paraventricular nucleus of the hypothalamus (PVH) is one of the pivotal wake-promoting nuclei ^5,6^, which has dense reciprocal connections with numerous neuroanatomical sites ^6^ that promote both wakefulness and emergence from inhaled GA, including the lateral septum (LS) ^7^, paraventricular thalamus (PVT) ^8,9^, and parabrachial nucleus (PB) ^10–14^. The PVH mainly consists of glutamatergic neurons ^15^. Specifically activating PVH glutamatergic neurons using chemogenetic approaches potently enhances the proportion of wakefulness, while lesions of PVH glutamatergic neurons has been shown to lead to serious hypersomnia in both mice and patients ^5,6^. As a subgroup of PVH glutamatergic neurons ^16^, corticotropin-releasing hormone neurons in the PVH (PVH^CRH^) neurons are involved in hypothalamic circuitry underlying stress-induced insomnia ^17^. Mild optogenetic stimulation of PVH^CRH^ neurons evoked persistent wakefulness, which mimicked insomnia elicited by restraint stress ^17^. Collectively, this evidence suggests that PVH^CRH^ neurons may play a role in regulating states of consciousness in GA.

The perioperative stress response is a spectrum of physiological changes occurring throughout different systems in the body, including neuroendocrine, metabolic, immunological, and hematological changes ^18^, as well as behavioral changes^19^. A prolonged stress response may stimulate excess cortisol release, immunosuppression, and an increase in proteolysis, which delays postoperative recovery and even leads to increased morbidity ^20,21^. As an important contributing factor of perioperative stress, GA-related stress has been neglected. Accumulated evidence has shown that the hypothalamic-pituitary-adrenal (HPA) axis is actively involved in the surgical stress during and after anesthesia ^22^. Post-operatively elevated cortisol secretion is considered a part of the surgical stress response ^23^. Distinct from other hypnotic agents, sevoflurane influences the endocrine response by increasing adrenocorticotrophic hormone and cortisol levels ^24,25^. Notably, PVH^CRH^ neurons serve as the initial node of the HPA axis, transducing the neuronal stress signal into a glucocorticoid secretion signal ^26^. Therefore, we hypothesize that PVH^CRH^ neurons may also act as a crucial node modulating the stress response of sevoflurane GA.

To test our hypothesis, we first performed global screening of active neurons during the post-anesthesia period in the mouse brain. We found that PVH^CRH^ neurons were preferentially active during the post-anesthesia period. After observing changes in calcium signals in PVH^CRH^ neurons during the induction and maintenance of, and recovery from sevoflurane GA along with a robust post-anesthesia stress response, we used chemogenetic and optogenetic manipulations to demonstrate how PVH^CRH^ neurons modulate sevoflurane GA. The results showed that the activation of PVH^CRH^ neurons conferred resistance to sevoflurane GA. Strikingly, inhibition of PVH^CRH^ neurons facilitated induction of sevoflurane GA and alleviated post-anesthesia stress responses. Hence, our findings uncover a vital brain site that modulates the state of consciousness as well as the stress response in sevoflurane GA.

## 2. Results

### 2.1. Identification of active neurons during the post-anesthesia period

Initially, to gain access to the active neurons during the post-anesthesia period, we subjected the mice to either sevoflurane-oxygen anesthesia (sevo) or oxygen exposure alone (control) for 30 min, followed by examining the whole-brain c-fos expression 3 h later (***Figure 1A***). Considering the time of the post-anesthesia stress response and the peak time of c-fos protein expression (2–3 h) after neuronal activation ^27^, a higher level of c-fos expression was observed in the PVH of sevo mice than that in control mice (***Figure 1B and C***). Increased staining was also observed in the anterior olfactory nucleus (AON), LS, ventromedial preoptic nucleus (VMPO), locus coeruleus (LC), and nucleus of the solitary tract (Sol) **(*Figure 1B, C and Figure 1-figure supplement 1A-C***). Interestingly, the PVH, traditionally considered a classic stress center, was the brain region with the highest relative expression in the sevo group ^28^. In contrast, other stress-related brain regions, such as the nucleus accumbens, the ventral tegmental area, and the central amygdaloid nucleus, did not exhibit robust c-fos expression (***Figure 1-figure supplement 2A-D***). Additionally, the medioventral periolivary nucleus (MVPO) showed decreased c-fos levels after inhalation of sevoflurane (***Figure 1B*** *and* ***Figure 1-figure supplement 1C***). To confirm specific markers for the c-fos-positive neurons in the PVH, we labelled PVH^CRH^ neurons by injecting a Cre-dependent adeno-associated virus (AAV) expressing mCherry into the PVH of CRH-Cre mice because CRH staining requires special pre-treatment (see Experimental Section, ***Figure 1-figure supplement 3***), and found that mCherry neurons shown high proportion of CRH immunoreactivity in the PVH (***Figure 1D top panel***). As shown by the colocalization of c-fos and CRH signals, approximately 65.71% of activated PVH neurons were CRH neurons, and approximately 73.91% of CRH neurons in the PVH were activated after exposure to sevoflurane GA (***Figure 1D bottom panel, E***). These results indicate that sevoflurane GA elicits robust activation of PVH^CRH^ neurons *in vivo*.

**Figure 1.**
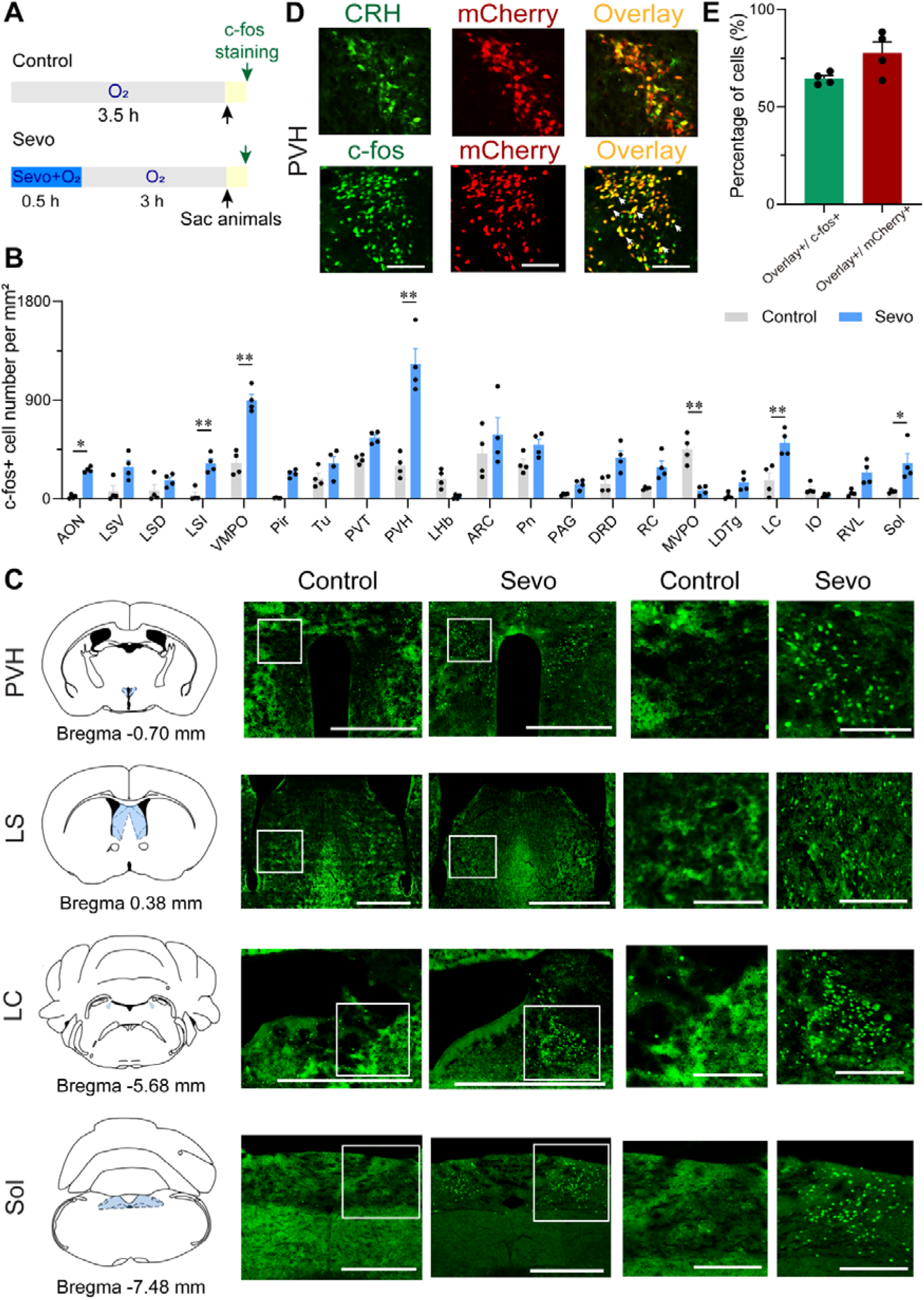
Whole-brain mapping of active neurons during the post-anesthesia period. **A.** Protocol for c-fos staining. Animals were exposed to sevoflurane or oxygen for 30 min and were sacrificed 3 h after the treatment. **B.** Quantification of the number of c-fos+ cells per mm^2^ in different brain regions (n = 4, unpaired two-tailed *t*-test, \**p* < 0.05, \*\**p* < 0.01). AON, anterior olfactory nucleus; ARC, arcuate hypothalamic nucleus; DRD, dorsal raphe nucleus, dorsal part; IO, inferior olive; LC, locus coeruleus; LDTg, laterodorsal tegmental nucleus; LHb, lateral habenular nucleus; LSD, lateral septal nucleus, dorsal part; LSI, lateral septal nucleus, intermediate part; LSV, lateral septal nucleus, ventral part; MVPO, medioventral periolivary nucleus; PAG, periaqueductal gray; Pir, piriform cortex; Pn, pontine nuclei; PVH, paraventricular nucleus of the hypothalamus; PVT, paraventricular thalamus; RC, raphe cap; RVL, rostroventrolateral reticular nucleus; Sol, nucleus of the solitary tract; Tu, olfactory tubercle; VMPO, ventromedial preoptic nucleus. **C.** Representative images of c-fos staining in the PVH, LS, LC, and Sol for a control and an experimental animal exposed to sevoflurane GA. Scale bar, 500 µm (left), 250 µm (right, enlarged). **D.** Representative images showing colocalization of CRH immunoreactivity and mCherry expression (upper panel, scale bar = 100 µm), c-fos and mCherry expression (bottom panel, scale bar = 100 µm) in the PVH of CRH-Cre mice. Arrowheads indicate co-labeled neurons. **E.** Quantification of the percentage of mCherry+ cells in the c-fos+ population (green) and the percentage of c-fos+ cells in the mCherry+ population (red) in the PVH.

### 2.2. Population activity of PVH^CRH^ neurons in response to sevoflurane GA

To reveal the real-time responses of PVH^CRH^ neurons to sevoflurane GA, we monitored the temporal dynamics of PVH^CRH^ neuronal activity in CRH-Cre mice during the whole process of sevoflurane GA (i.e., induction, maintenance, emergence, and recovery phases) using *in vivo* fiber photometry. A Cre-dependent AAV expressing the fluorescent calcium indicator, jGCaMP7b (AAV2/9-hSyn-DIO-jGCaMP7b-WPRE-pA), was injected into the PVH of CRH-Cre mice (***Figure 2A***). The expression of the virus is shown in ***Figure 2B***. During the whole experimental process, the mice were placed in a vertical cylindrical cage for sevoflurane GA (***Figure 2C***). As shown in ***Figure 2D***, the neural activity declined with exposure to sevoflurane and increased with the cessation of sevoflurane inhalation. In detail, compared to the awake baseline, PVH^CRH^ neurons showed significantly decreased neuronal activity during the 30-min exposure to 1.6% sevoflurane. By analyzing the Ca^2+^ signals over four periods: pre (–30 to 0 min), during (anesthesia period, 0 to 30 min), post 1 (30 min to time to recovery of righting reflex [RORR]), and post 2 (RORR to baseline), we found that the population activities of PVH^CRH^ neurons were significantly blunted by sevoflurane exposure (during versus pre: –31.11% ± 9.11%; *p* = 0.027) and gradually returned during the recovery period (post 1 versus during: 5.84% ± 1.19%; *p* = 0.651). These findings suggest that the activities of PVH^CRH^ neurons were altered across distinct concentrations of sevoflurane.

**Figure 2.**
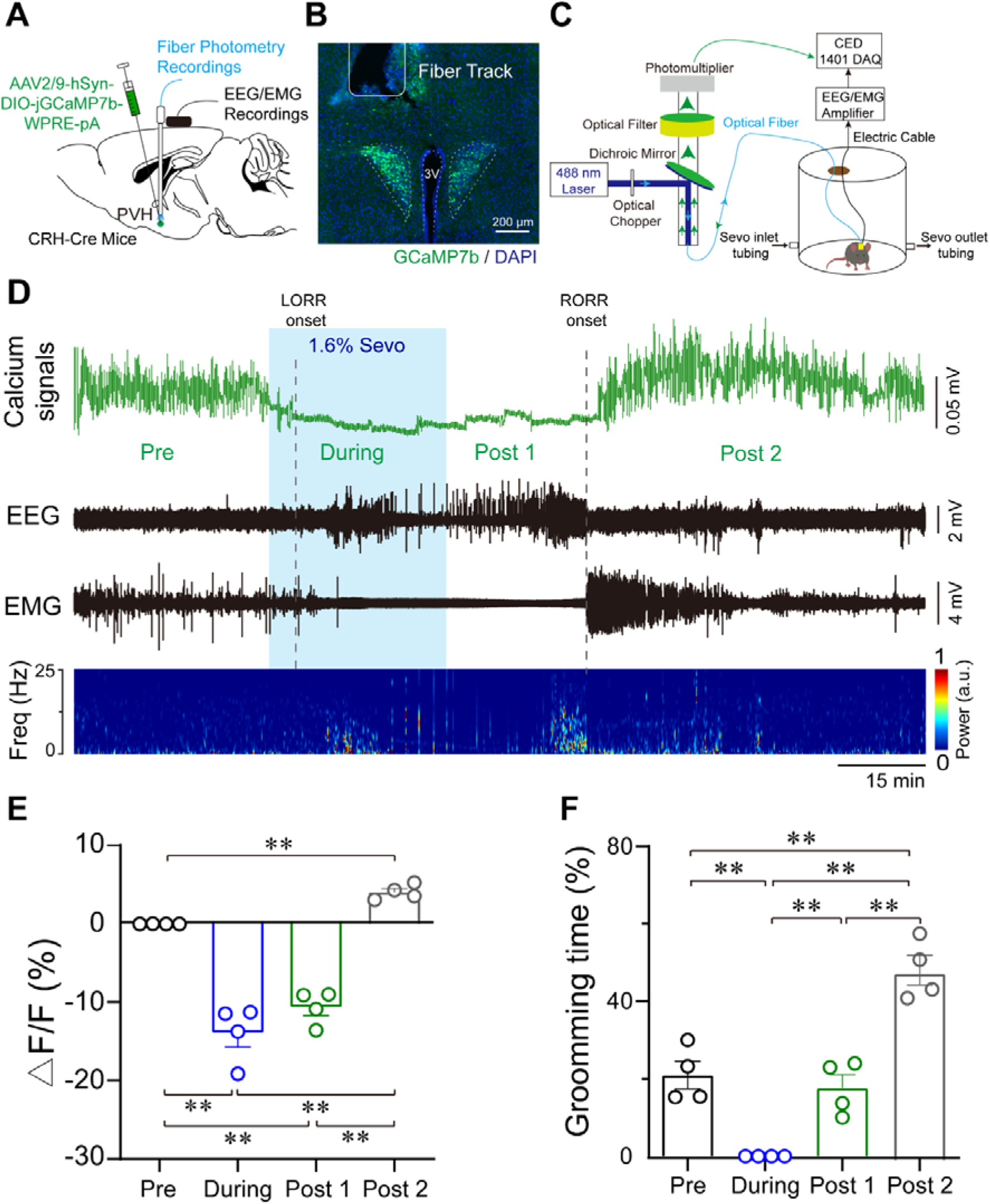
Population activities of PVH^CRH^ neurons in response to sevoflurane GA. **A.** Diagram of the virus injection, EEG/EMG electrode, and optic fiber implantation sites of CRH-Cre mice. **B.** jGCaMP7b/DAPI immunofluorescence in CRH neurons and track of the optic fiber implanted above the PVH; scale bar, 200 μm. Viral expression of jGCaMP7b and placement of the fiber-optic probe above the PVH. **C.** Schematic of the recording configuration. **D-E.** Time courses of Ca^2+^ signals following sevoflurane anesthesia **D** and quantification of Ca^2+^ signal changes before, during, and after (post 1 and post 2 periods) sevoflurane inhalation (**E**, n = 4, F (3, 12) = 61.49, *p* < 0.001, one-way ANOVA with Tukey’s post-hoc test; pre vs. during, pre vs. post 1, during vs. post 2, post 1 vs. post 2, *p*\*\*< 0.001; pre vs. post 2, *p* = 0.0894; during vs. post 1, *p* = 0.2012). Freq, frequency; LORR, loss of right reflexing; RORR, recovery of right reflexing; Sevo, sevoflurane. **F.** Time percentage of self-grooming before, during, and after (post 1 and post 2 periods) sevoflurane inhalation (n = 4, F (3, 12) = 76.87, *p* < 0.001, one-way repeated measures ANOVA with Tukey’s post-hoc test; pre vs. during, pre vs. post 1, *p* = 0.0005; pre vs. post 2 during vs. post 2, post 1 vs. post 2, *p*\*\* <0.0001; during vs. post 1, *p* > 0.9999).

Notably, PVH^CRH^ neurons exhibited higher Ca^2+^ signals compared to the awake baseline after a 30-min cessation of sevoflurane inhalation. After comparing the Ca^2+^ signals of the awake baseline period (pre) and post-firing period (post 2), we found that the population activities of PVH^CRH^ neurons increased potently after 1.6% sevoflurane exposure (***Figure 2E***, pre versus post 2: 40.38% ± 4.75%; *p* = 0.002). Interestingly, this hyperactivity phenomenon, lasting approximately 1 h before returning to baseline levels, was accompanied by enhanced self-grooming (***Figure 2F***), which is considered a stress-related behavior ^29^. These results prompted us to speculate that the hyperactivities of PVH^CRH^ neurons may play a vital role in the post-anesthesia stress response.

### 2.3. Characterizations of sevoflurane GA-induced grooming

Stress in animals often results in grooming and other repetitive behaviors ^30^. As these displacement activities are believed to have adaptive values to stress ^31,32^, we employed excessive self-grooming as an indicator of stress response of sevoflurane GA. Next, we interrogated the characteristics of sevoflurane GA-induced grooming.

Considering that grooming behaviors are context-sensitive and could be reflected in their sequence patterns or microstructure, we analyzed these characteristics of sevoflurane GA-induced grooming. Following the termination of sevoflurane GA, mice stayed awake for approximately 1.5 h, during which they exhibited robust self-grooming with increased duration per bout and total time spent grooming (***Figure 3-figure supplement 4A***). Because of the consistent trend, we divided the post-anesthesia period (post 2) into two sections (0–50 min versus 50–100 min). Compared to the former period, the latter period showed higher percentages of incorrect phase transition and interrupted bouts (see ‘Experimental Section’), indicating elevated stress levels (***Figure 3A and B***); thus, we selected the latter as the representative period to investigate sevoflurane GA-induced grooming.

**Figure 3.**
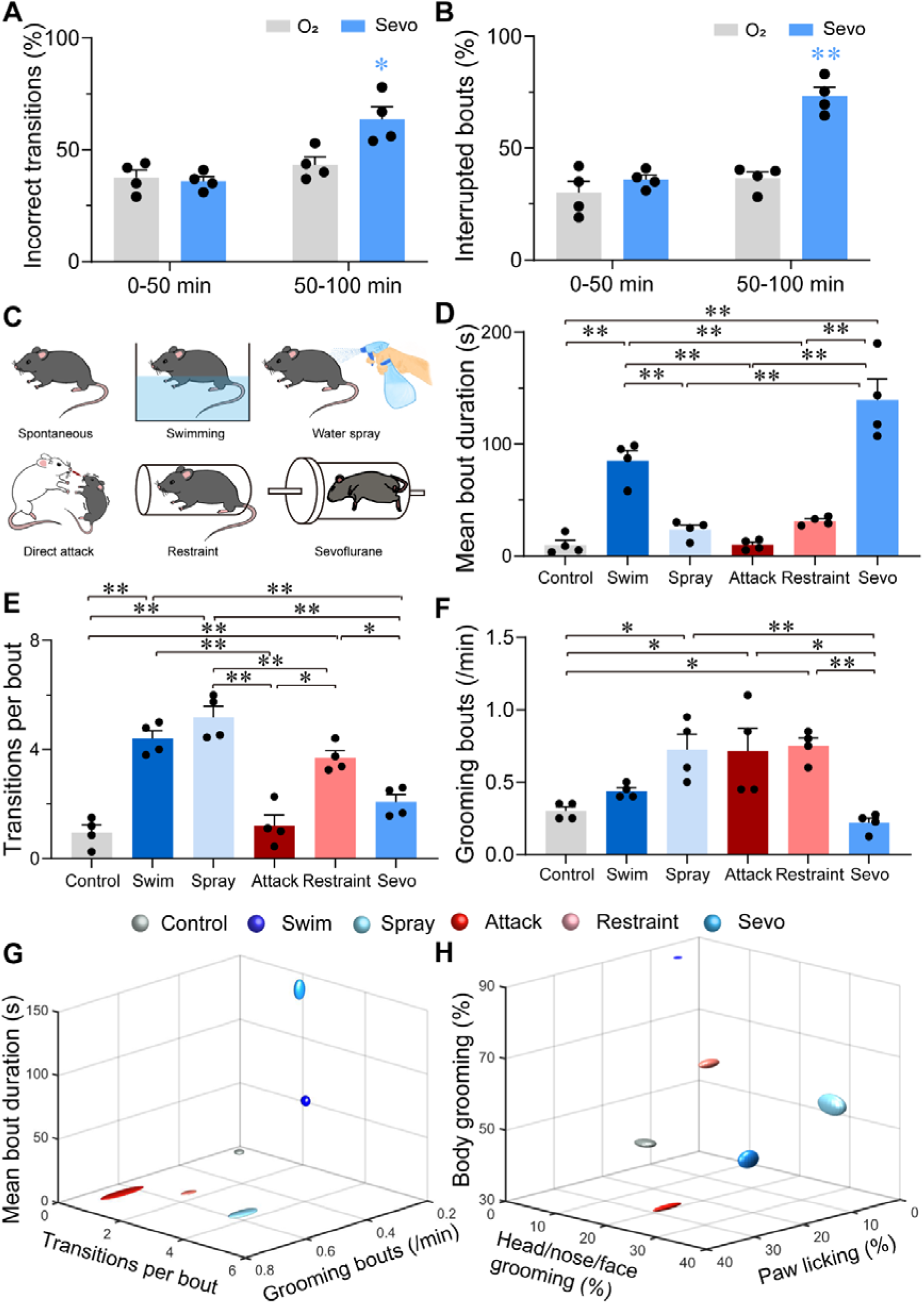
Characterizations of sevoflurane GA-induced grooming. **A.** Quantification of incorrect transitions with respect to the cephalocaudal sequence of stereotypic grooming patterns in the post-anesthesia period (n = 4, two-way ANOVAs followed by Sidak’s test, F (1, 6) = 8.646, *p* = 0.0259, 0–50 min: O_2_ vs sevo, *p* = 0.9545, t = 0.2767, df = 12, 95% CI = -12.34 to 15.34; 50–100 min, O_2_ vs sevo, *p* = 0.0055, t = 3.753, df = 12, 95% CI = -34.19 to -6.504). **B.** Number of interrupted bouts in the post-anesthesia period (n = 4, two-way ANOVAs followed by Sidak’s test, F (1, 6) = 19.14, *p* = 0.0047, 0–50 min: O_2_ vs sevo, *p* = 0.4770, t = 1.139, df = 12, 95% CI = -19.45 to 7.446; 50–100 min, O_2_ vs sevo, \*\**p* < 0.01, t = 6.969, df = 12, 95% CI = -50.15 to -23.26). **C.** Six representative grooming models, including spontaneous (control), swimming, water spray, physical attack, body restraint, and sevoflurane GA-induced grooming. **D-F.** The mean bout duration (**D**, F (5, 18) = 34.52, *p* < 0.0001), transitions per bout (**E**, F (5, 18) = 30.32, *p* < 0.0001), and grooming frequency (bouts per min, **F**, F (5, 18) = 7.935, *p* =0.0004) varied across the models (n = 4, one-way ANOVA with Tukey’s post-hoc test. \**p* < 0.05; \*\**p* < 0.01). **G-H.** 3-D plot of bout frequency, bout duration, and transitions per bout (**G**); the percentage of time spent grooming different body parts (**H**). The dimension of the symbol along an axis is defined by the SD of the corresponding parameter.

Subsequently, we compared sevoflurane GA-induced grooming with four grooming models related to physical and emotional stress, including those elicited by free swimming, water spray, physical attack, and body restraint (***Figure 3C***). The former two models induce more physical stress by moistening the fur, while the latter two models are often considered models for emotional stress ^33^. In line with previous reports ^31,32,34,35^, the mice in these two emotional stress-related groups spent a higher percentage of time paw licking, while those in the two fur moistening groups spent most time in grooming the body (***Figure 3-figure supplement 4D*** *and **E***). Interestingly, for transition bouts and grooming bouts per min, sevoflurane GA-induced grooming during 50–100 min was close to spontaneous grooming (***Figure 3E-G***), but it shared no similarities with these four grooming models in terms of the mean bout duration (***Figure 3D and G***). A plot of the number of times spent grooming different body regions also showed no similarity among sevoflurane GA-induced grooming and other models (***Figure 3H and Figure 3-figure supplement 4D-F***). The combination of these parameters suggested a low level of stress at the beginning of the post-anesthesia period followed by a higher and unique level of stress with different parameters (longest mean bout duration, lowest bout frequency, and average distribution of grooming time spent grooming different body parts).

### 2.4. Modulation of PVH neurons altered the induction and emergence of sevoflurane GA

To investigate the role of PVH^CRH^ neurons during sevoflurane GA, we micro-injected AAV-DIO-hM3Dq-mCherry or AAV-DIO-hM4Di-mCherry or AAV-DIO-mCherry into the bilateral PVH of CRH-Cre mice, respectively (***Figure 4A***). The c-fos expression overlaid with hM3Dq or hM4Di is shown in ***Figure 4B***. We then performed behavioral experiments to determine the effect of chemogenetic activation or inhibition of PVH^CRH^ neurons on the sensitivity, induction, and emergence of sevoflurane GA. First, the sensitivity of sevoflurane GA was evaluated by a minimum inhaled sevoflurane concentration that evoked loss of righting reflex (LORR), which is a potent behavioral indicator of the onset of GA. We found that PVH^CRH^ neuron-activated mice were resistant to higher concentrations of sevoflurane than the vehicle group (***Figure 4D***), whereas the PVH^CRH^ neuron-inhibited mice were more vulnerable to sevoflurane GA (***Figure 4E***). The right-shift of the dose-response curve along with the augmentation of the 50% effective concentration (EC_50_) in the PVH^CRH^ neuron-activated mice suggested that PVH^CRH^ neuron activation lowered sevoflurane sensitivity during sevoflurane induction. Conversely, the left-shift of the dose-response curve combined with the decrease in EC_50_ in the PVH^CRH^ neuron-inhibited mice revealed higher sevoflurane sensitivity during induction. According to the dose-response curves (***Figure 4-table supplement 1***), the EC_50_ of sevoflurane in term of LORR from the PVH^CRH^ neuron-activated mice (n = 10) was 1.650% versus 1.516% of the vehicle group, and the EC_50_ of sevoflurane in term of RORR also increased (***Figure 4-figure supplement 5A***). As for the chemogenetic inhibition experiments, the EC_50_ of the PVH^CRH^ neuron-inhibited mice (n = 8) was 1.337% versus 1.500% of the vehicle mice, and the EC_50_ of sevoflurane in term of RORR also reduced (***Figure 4-figure supplement 5B***). The EC_50_ of sevoflurane in term of either RORR or LORR did not show significant change in control group (***Figure 4-figure supplement 5C*** *and **D***).

**Figure 4.**
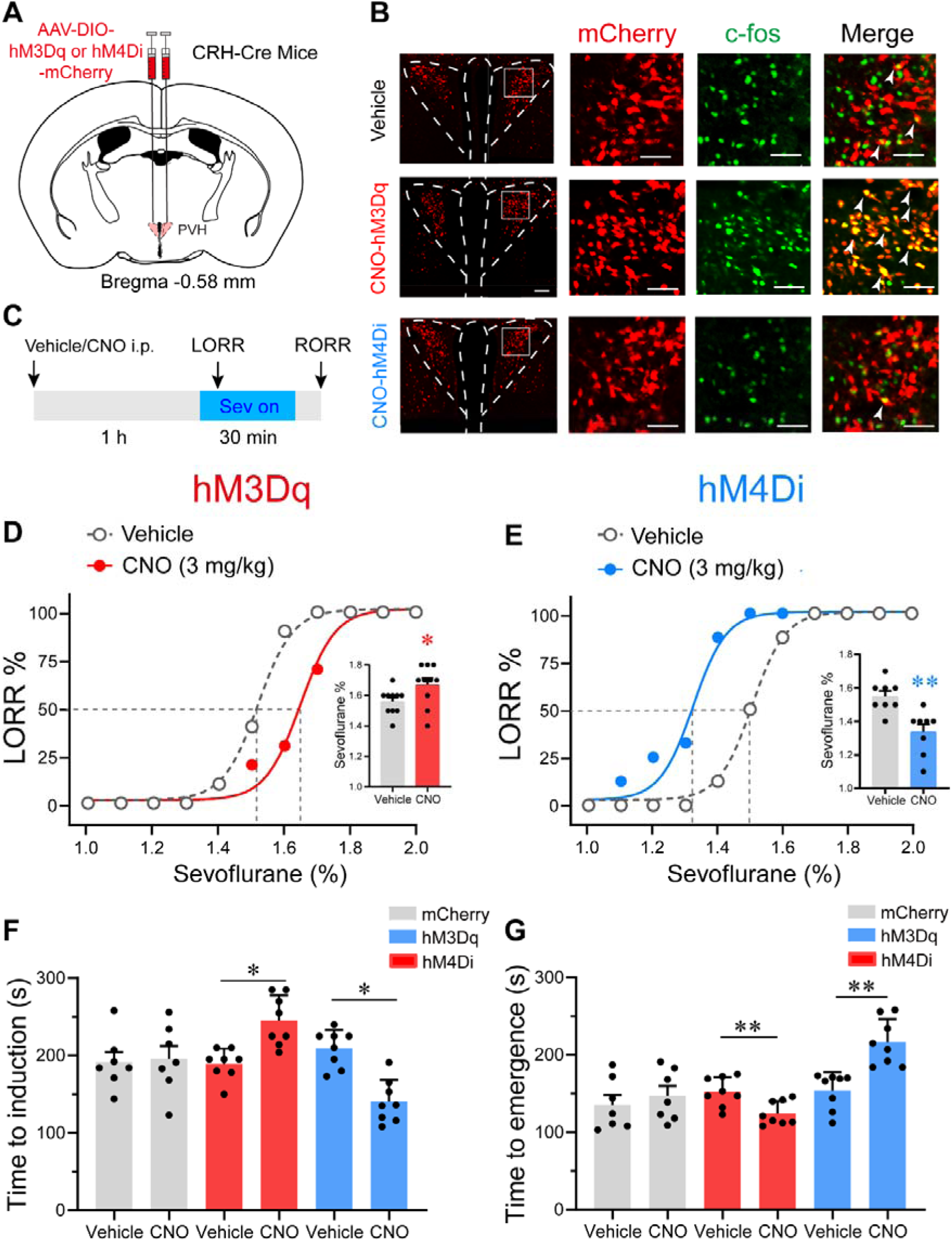
Chemogenetic modulation of PVH^CRH^ neurons bidirectionally altered induction of and emergence from sevoflurane GA. **A.** Schematic of AAV-DIO-hM3Dq-mCherry or AAV-DIO-hM4Di-mCherry or AAV-DIO-mCherry injected into the PVH of CRH-Cre mice. **B.** Left: representative images of mCherry/c-fos immunofluorescence in CRH neurons after vehicle or CNO treatment; scale bars, 200 μm. Right: magnified images are shown; scale bar, 200 μm. Arrowheads indicate co-labeled neurons. **C.** Timelines of sevoflurane anesthesia-related behavioral tests measuring induction time (LORR) and emergence time (RORR). **D-E.** Dose-response curves showing the percentages of mice exhibiting LORR in response to incremental sevoflurane concentrations for the vehicle and CNO groups. Inset: the sevoflurane concentrations at which each mouse exhibited LORR are shown (**D**, hM3Dq group, n = 10, paired *t*-test, *p* = 0.0411, t = 4.714, df = 9, 95% CI = 0.05 to 0.16; **E**, hM4Di group, n = 8, paired *t*-test, *p* = 0.0021, t = 9.375, df = 7, 95% CI = -0.266 to -0.1589). **F.** Induction time with 2% sevoflurane exposure after intraperitoneal injections of vehicle or CNO for 1 h (mCherry group, n = 7, paired *t*-test, *p* = 0.847, t = 0.2014, df = 6, 95% CI = -47.77 to 56.35; hM3Dq group, n = 8, paired *t*-test, *p* = 0.0498, t = 5.545, df =7, 95% CI = 32.26 to 80.24; hM4Di group, n = 8, paired *t*-test, *p* = 0.0060, t = 4.573, df =7, 95% CI = -104.1 to -33.14). **G.** Emergence time with 2% sevoflurane exposure for 1 h after intraperitoneal injections of vehicle or CNO (mCherry group, n = 7, paired *t*-test, *p* = 0.8298, t = 0.9286, df =6, 95% CI = -19.15 to 42.58; hM3Dq group, n = 8, paired *t*-test, *p* = 0.0048, t = 4.057, df =7, 95% CI = -43.72 to -11.53; hM4Di group, n = 8, paired *t*-test, *p* = 0.0057, t = 3.922, df =7, 95% CI = 24.77 to 99.98).

We then measured the induction time (time to LORR) and emergence time (RORR) following the experiment procedure outlined in ***Figure 4C***. In the chemogenetic activation experiments, clozapine-N-oxide (CNO) pre-treatment led to increased induction time after 2% sevoflurane exposure (***Figure 4F***). Concomitantly, the PVH^CRH^ neuron-activated mice demonstrated faster emergence from 2% sevoflurane GA (***Figure 4G***). As for the chemogenetic inhibition manipulation, an obvious shortened induction time and a prolonged emergence time were observed after CNO pre-treatment (***Figure 4H and I***). These results further imply that chemogenetically activating PVH^CRH^ neurons increased resilience to sevoflurane and promoted awakening from sevoflurane GA, whereas chemogenetic inhibition manipulation of PVH^CRH^ neurons exerted contrary effects. To further examine the role of PVH^CRH^ neurons in sevoflurane GA, we genetic ablated PVH^CRH^ neurons (***Figure 4-figure supplement 6A*** *and **B***) and found similar effects with inhibition group in terms of dose-response curve. The EC_50_ of sevoflurane in terms of LORR decreased from 1.491 to 1.358% compared with control group (***Figure 4-figure supplement 6C***), and lesion of PVH^CRH^ neurons decreased EC_50_ of sevoflurane on RORR from 1.420 to 1.326% (***Figure 4-figure supplement 6D***). Meanwhile, genetic ablation of PVH^CRH^ neurons significantly facilitated induction and slowed emergence of sevoflurane GA (***Figure 4-figure supplement 6E and F***), which show lower induction time and longer emergence time than the inhibition group (***Figure 4D and E***).

Considering the temporal limitation of chemogenetic manipulation, we next employed optogenetic methods with millisecond-scale control of neuronal activities to verify the role of PVH^CRH^ neurons in controlling sevoflurane GA. To this end, we stereotaxically injected AAVs expressing channelrhodopsin-2 (AAV2/9-hEF1a-DIO-hChR2-mCherry-pA) or AAV-DIO-mCherry into the bilateral PVH of CRH-Cre mice with optical fibers implanted (***Figure 4-figure supplement 7A***). We verified the expression of ChR2 and the optical fiber locations after behavior experiments (***Figure 4-figure supplement 7B***). Mice were placed in the same vertical cylindrical cage as the apparatus for *in vivo* fiber photometry for sevoflurane GA. Optical blue-light stimulation (5 ms, 30 Hz, 20–30 mW/mm^2^) was applied at the onset of induction and ended until LORR or started at the end of sevoflurane inhalation and continued until RORR (***Figure 4-figure supplement 7C***). We found that optical activation of PVH^CRH^ neurons prominently delayed the induction process (***Figure 4-figure supplement 7D***) and accelerated the recovery process (***Figure 4-figure supplement 7E***) compared to the mCherry control, indicating that optical activation of PVH^CRH^ neurons could prompt the emergence from sevoflurane GA.

### 2.5. Optogenetic stimulation of PVH neurons induced rapid emergence from steady sevoflurane GA state

We next applied 60 s of optical blue-light stimulation (5 ms, 30 Hz, 20–30 mW/mm^2^) after mice had maintained LORR for 30 min during steady GA state maintained by 2% sevoflurane. At this stage, the electroencephalogram (EEG) patterns of the mice were characterized by an increase in low-frequency, high-amplitude activity, which resembles the EEG pattern of non-rapid eye movement sleep (slow-wave sleep). The optogenetics experiments were performed in the similar apparatus with partially sealed chamber in the ***Figure 2C***. Photostimulation of PVH^CRH^ neurons at 30 Hz during this state reliably elicited a brain-state transition from slow-wave activity to a low-amplitude, high-frequency activity in ChR2 mice, but not in mCherry mice (***Figure 5A and B***). We also observed behavioral emergence (including body movements of limbs, head, and tail, righting, and walking) (***Table S2, Supporting Information***) combined with enhanced electromyography (EMG) activity after photostimulation at 30 Hz. Spectral analysis of EEG data revealed that acute photostimulation of PVH^CRH^ neurons at 30 Hz elicited a prominent decrease in delta power (60.83% ± 1.35% versus 40.23% ± 5.30%; *p* = 0.045, t = 7.406, df = 40, 95% CI = 12.46 to 26) and an increase in alpha power (7.47% ± 0.41% versus 12.42% ± 0.93%; *p* = 0.033, t = 1.721, df = 40, 95% CI = -11.24 to 2.303) and beta power (6.01% ± 0.245% versus 11.23% ± 1.27%; *p* = 0.025, t = 1.727, df = 40, 95% CI = -11.25 to 2.286; ***Figure 5D and F***). However, there was no significant difference between the EEG spectrum of pre-stimulation and stimulation periods in mCherry mice (***Figure 5C and E***). These findings revealed that optogenetic activation of PVH^CRH^ neurons was sufficient to induce cortical activation and behavioral emergence during the steady sevoflurane GA state.

**Figure 5.**
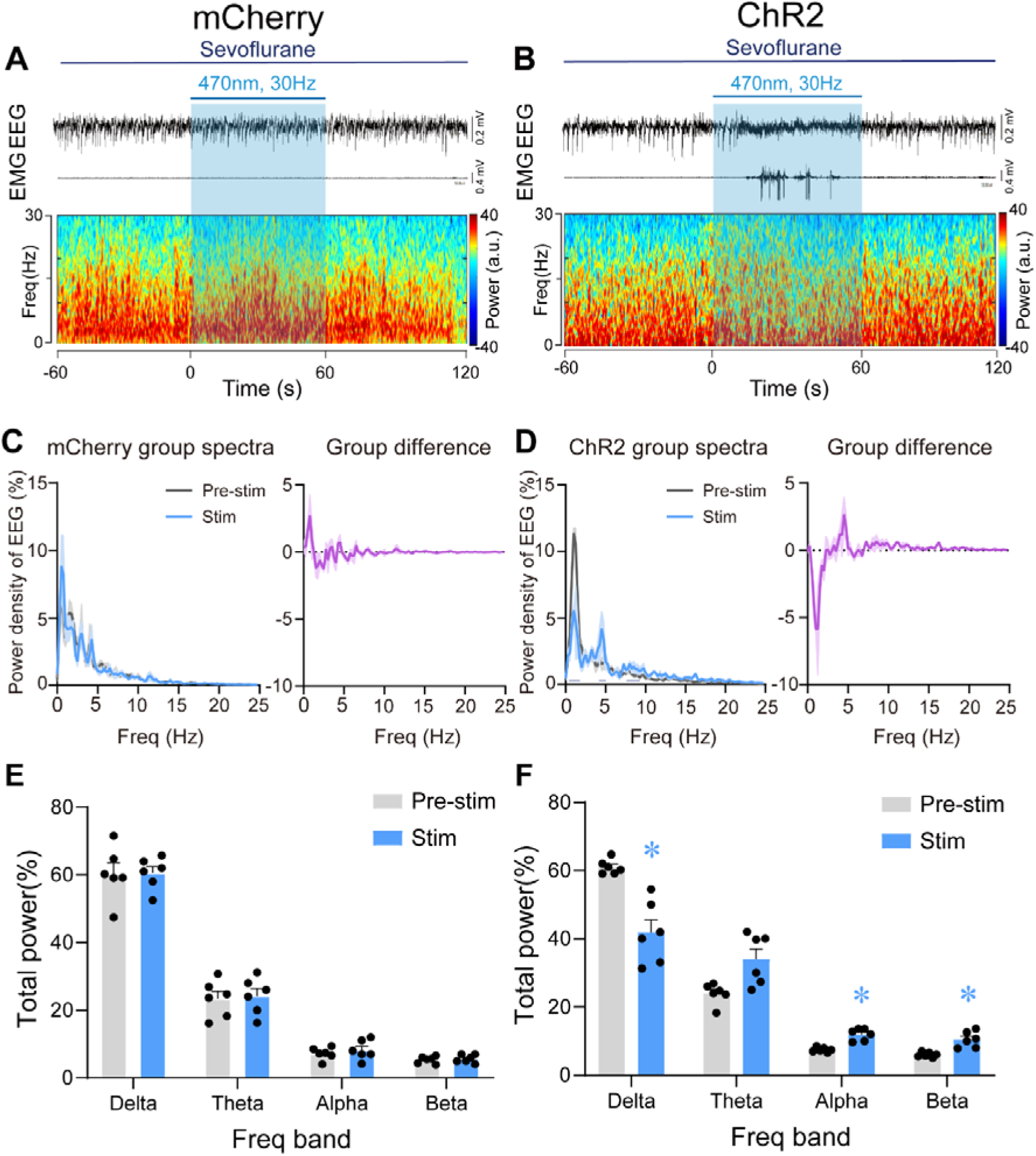
Optogenetic stimulation of PVH^CRH^ neurons induced cortical activation and behavioral emergence during continuous steady-state sevoflurane GA. **A-B.** Typical examples of EEG, EMG, and EEG power spectral in a mouse injected with AAV-DIO-mCherry(**A**) or AAV-DIO-ChR2-mCherry (**B**) following acute photostimulation (30 Hz, 5 ms, 60 s) during continuous steady-state sevoflurane GA. Time 0 indicates the beginning of photostimulation. The blue shadow indicates the 60-s duration of blue light stimulation. **C-D.** Left: normalized group power spectral densities from PVH^CRH^-mCherry (**C**) or PVH^CRH^-ChR2 (**D**) mice with pre photostimulation (gray) and photostimulation (blue). Dark blue lines in d indicate the power band with significant difference. Right: differences between normalized group power spectral densities from PVH^CRH^-mCherry (**C**) or PVH^CRH^-ChR2 (**D**). **E-F.** Power percentage changes in cortical EEG before (gray) and during (blue) photostimulation at 30 Hz in PVH^CRH^-mCherry (**E**) or PVH^CRH^-ChR2 mice (**F**) during continuous steady-state sevoflurane GA (n = 6, two-way ANOVAs followed by Sidak’s test, \**p* < 0.05).

### 2.6. Optogenetic stimulation of PVH^CRH^ neurons induced EEG activation during burst-suppression oscillations induced by deep sevoflurane GA

We further explored the role of PVH^CRH^ neurons in the regulation of deep sevoflurane GA, during the burst-suppression phase, in which the EEG pattern is characterized by flat periods interspersed with periods of alpha and beta activity ^36^. After the maintenance of burst-suppression EEG mode for at least 5 min, we delivered 60 s of optical stimulation (5 ms, 30 Hz, 20–30 mW/mm^2^) and found that EEG activation mainly occurred with an average latency of 20 s and maintained for a few minutes before returning to burst-suppression mode. However, the EMG activity was not significantly different between pre-/post-stimulation periods of mCherry mice (***Figure 6A*** *and* ***Figure 6-figure supplement 8A***) and ChR2 mice (***Figure 6B*** *and* ***Figure 6-figure supplement 8B***). Statistical analysis of 1 min of EEG recordings (optical stimulation for 20 s and 20 s before and after photostimulation) demonstrated a robust decline in delta power (from 53.85% ± 2.58% to 41.46% ± 2.37%, *p* = 0.005, df = 60, 95% CI = 12.3 to 22.34) and an increase in theta power (from 24.25% ± 3.50 to 32.07% ± 2.12%, *p* = 0.017, df = 60, 95% CI = -15.74 to -5.699), alpha power (from 6.72% ± 1.16% to 11.15% ± 1.37%, *p* = 0.0062, df = 60, 95% CI = -11.69 to -1.652), and beta power (from 4.00% ± 0.55% to 7.56% ± 0.01%, *p* = 0.0317, df = 60, 95% CI = -10.43 to -0.394; ***Figure 6D and G***). As for the mCherry mice, 60 s of blue-light stimulation had no significant effect on either EEG or EMG activity (***Figure 6A, C and E***). Photostimulation of PVH^CRH^ neurons in ChR2 mice also caused a significant reduction in BSR (Pre-stim vs. Post-stim, *p* < 0.01, df = 15, 95% CI = 23.53 to 48.14; Pre-stim vs. Stim, *p* < 0.01, df = 15, 95% CI = 17.36 to 41.97; ***Figure 6H***). In contrast, there were no significant changes in mCherry mice with 30 Hz photostimulation (***Figure 6F***). These results collectively suggested that PVH^CRH^ neuronal activation was sufficient to facilitate EEG cortical activation but not to promote behavioral emergence during deep states of sevoflurane GA with burst-suppression oscillations.

**Figure 6.**
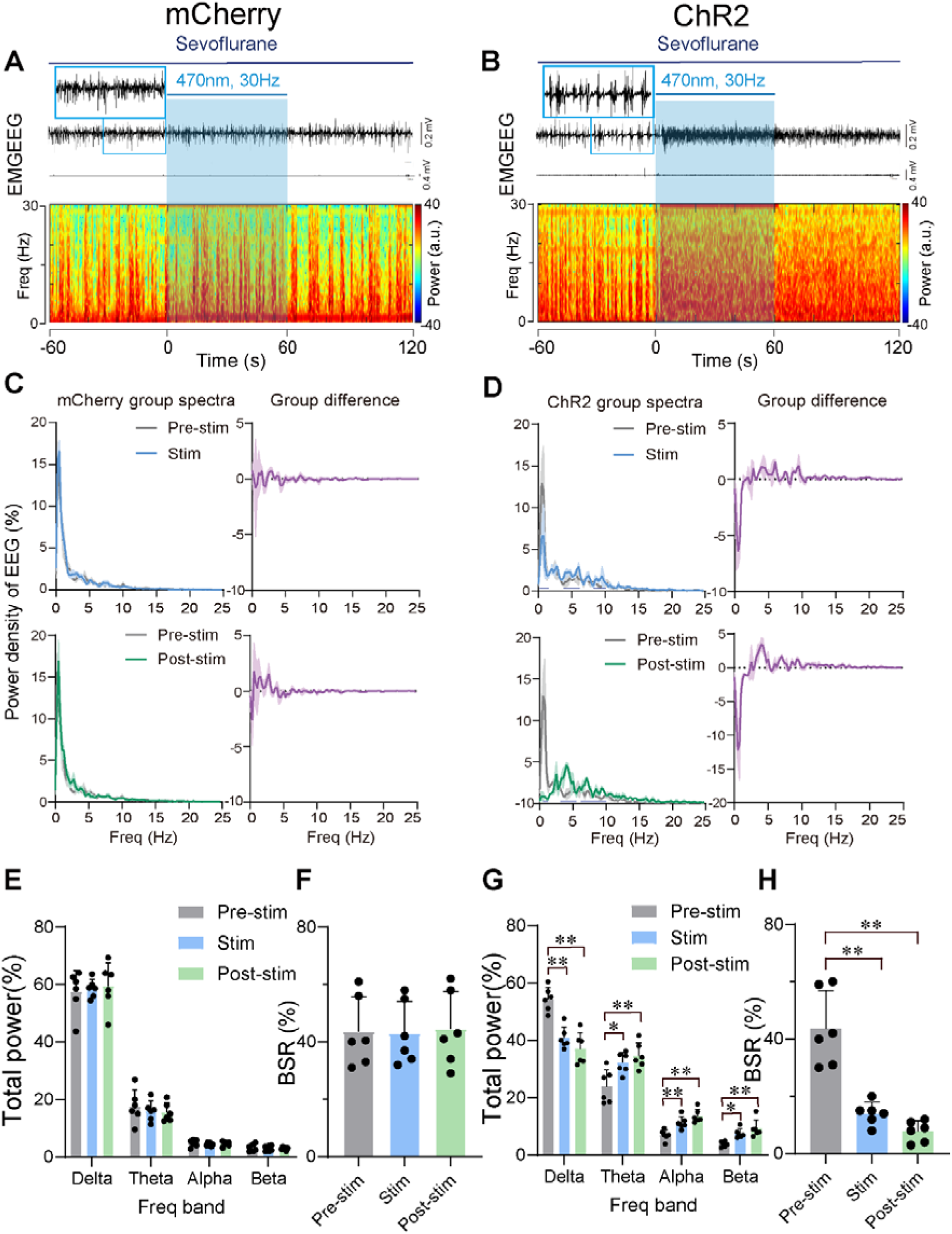
Optogenetic stimulation of PVH^CRH^ neurons induced cortical activation during burst-suppression oscillations induced by deep sevoflurane GA. **A-B.** Typical examples of EEG, EMG, and EEG power spectral in a mouse injected with AAV-DIO-mCherry (**A**) or AAV-DIO-ChR2-mCherry (**B**) following acute photostimulation (30 Hz, 5 ms, 60 s) during burst-suppression oscillations. Time 0 indicates the beginning of photostimulation. The blue shadow indicates the 60-s duration of blue light stimulation. **C-D.** Top left: normalized group power spectral densities from PVH^CRH^-mCherry mice (**C**) and PVH^CRH^-ChR2 (**D**) with pre photostimulation (gray) and photostimulation (blue); Top right: differences between normalized group power spectral densities from PVH^CRH^-mCherry mice (**C**) and PVH^CRH^-ChR2 mice (**D**). Bottom left: normalized group power spectral densities from PVH^CRH^-mCherry mice (**C**) and PVH^CRH^-ChR2 mice (**D**) with pre photostimulation (gray) and post photostimulation (green); bottom right: differences between normalized group power spectral densities from PVH^CRH^-mCherry mice (**C**) and PVH^CRH^-ChR2 mice (**D**). Dark blue lines in panel d indicate the power band with significant difference. **E, G.** Power percentage changes in cortical EEG before (gray), during (blue), and post (green) photostimulation in PVH^CRH^-mCherry (**E**) or PVH^CRH^-ChR2 (**G**) mice during burst-suppression oscillations. **F, H.** BSR change before (gray), during (blue), and post (green) photostimulation in PVH^CRH^-mCherry (**G**) or PVH^CRH^-ChR2 (**H**) mice during burst-suppression oscillations (n = 6, two-way ANOVAs followed by Sidak’s test, \**p* < 0.05, \*\**p* < 0.01; Pre-stim vs. Stim, p < 0.01, df = 15, 95% CI = 17.36 to 41.97; Pre-stim vs. Post-stim, p < 0.01, df = 15, 95% CI = 23.53 to 48.14). Stim, stimulation; Freq, frequency.

### 2.7. Chemogenetic inhibition of PVH^CRH^ neurons compromised the stress response after sevoflurane GA

Given the hyperactivity of PVH^CRH^ neurons and accompanying increase in self-grooming after exposure to sevoflurane GA, we investigated whether PVH^CRH^ neurons are recruited while mice are subjected to post-anesthesia stress. We assessed the behavioral effect of chemogenetic inactivation of these neurons via the bilateral expression of the inhibitory hM4Di receptors (***Figure 7A***). Following the protocol shown in ***Figure 7B***, we found that inhalation of 1.6% sevoflurane induced higher expression of c-fos in the PVH, which was significantly suppressed by chemogenetic inhibition of PVH^CRH^ neurons (***Figure 7C***). Hypothalamic CRH neurons control the circulating levels of corticosteroid stress hormones in the body ^37^. In line with the c-fos expression, serum CRH levels showed similar changes after sevoflurane inhalation both with and without chemogenetic inhibition (***Figure 7D***), suggesting that peripheral CRH levels changed with the same direction of PVH^CRH^ neuronal activity.

**Figure 7.**
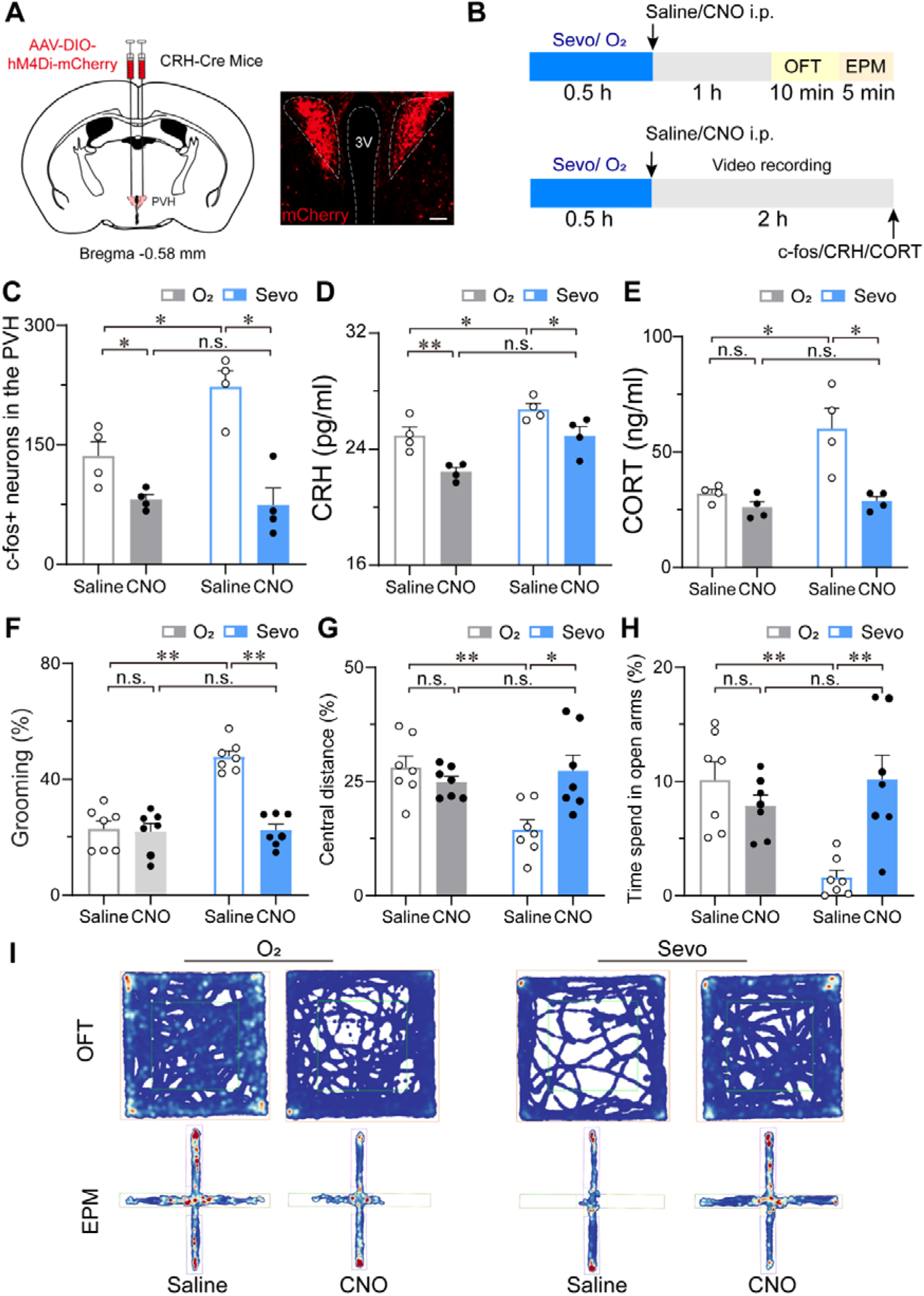
Chemogenetic inhibition of PVH^CRH^ neurons alleviated the stress response after sevoflurane GA. **A.** Schedule of sacrificing mice for c-fos quantification, collecting blood samples from mice subjected to sevoflurane or pure oxygen for CRH, and CORT measurement and behavior testing (OFT and EPM). **B.** The schedule of the video recording after inhalation of sevoflurane or pure oxygen for 30 min. **C.** Quantification of the number of c-fos-positive neurons in the PVH (n = 4, saline: O_2_ vs. sevo, *p* = 0.0017, t = 3.523, df = 12, 95% CI = -164.6 to -9.424; CNO: O_2_ vs. sevo, *p* = 0.7122, t = 0.2632, df = 12, 95% CI = -71.08 to 84.08). **D-E.** Serum CRH (**D**, n = 4, saline: O_2_ vs. sevo, *p* = 0.0456, t = 2.604, df = 12, 95% CI = -3.585 to -0.03; CNO: O_2_ vs. sevo, *p* = 0.0077, t = 3.567, df = 12, 95% CI = -4.255 to -0.705) and CORT (**E**, n = 4, saline: O_2_ vs. sevo, *p* = 0.0032, t = 4.059, df = 12, 95% CI = -44.73 to -10.19; CNO: saline: O_2_ vs. sevo, *p* = 0.9066, t = 0.4024, df = 12, 95% CI = -19.99 to 14.55) levels following the protocol in (a). **F-H.** Time percentage of self-grooming (**F**, n = 7, saline: O_2_ vs. sevo, *p* < 0.001, t = 7.23, df = 24, 95% CI = -33.26 to -16.76,; CNO: O_2_ vs. sevo, *p* = 0.9915, t = 0.1171, df = 24, 95% CI = -8.656 to 7.846), moving distances in the central areas of OFT (**G**, n = 7, saline (: O_2_ vs. sevo, *p =* 0.0014, t = 3.892, df = 24, 95% CI = 5.269 to 21.95; CNO: O_2_ vs. sevo, *p* = 0.7291, t = 0.7183, df = 24, 95% CI = -10.85 to 5.829) and time percentage of staying in the open arms of the EPM (**H**, n = 7, saline: O_2_ vs. sevo, *p =* 0.0007, t = 4.166, df = 24, 95% CI = 3.649 to 13.42; CNO: O_2_ vs. sevo, *p =* 0.4626, t = 1.137, df = 24, 95% CI = -7.217 to 2.559) after inhalation of sevoflurane or pure oxygen and administration of saline or CNO. Statistical comparisons were conducted using two-way ANOVA followed by Sidak’s tests. \**p* < 0.05, \*\**p* < 0.01, n.s., no significant differences). **I.** Representative heatmaps of OFT and EPM after inhalation of sevoflurane or pure oxygen and administration of saline or CNO.

Considering that corticosterone (CORT) is a hormone commonly used to measure stress in rodents ^38^, we measured the serum CORT levels and found that 30 min of 1.6% sevoflurane inhalation prominently evoked increased serum CORT concentrations after 1-h termination of sevoflurane compared to that in groups inhaling pure oxygen (both groups injected saline or CNO), which were restored to normal levels (***Figure 7E***). However, although the serum CRH levels of pure oxygen groups decreased after CNO injection, there was no significant difference in serum CORT concentrations between saline and CNO injection groups after inhalation of pure oxygen. These results implicate that the inhibition of PVH^CRH^ neurons lowers the high levels of serum CORT due to sevoflurane exposure.

Next, we observed the stress-related behavior after inhaling pure oxygen (O_2_) or 1.6% sevoflurane (sevo) for 30 min (***Figure 7B***). Analysis of self-grooming behavior (1.5 h after sevoflurane GA cessation) showed that the percentage of time spent self-grooming increased after 1.6% sevoflurane exposure (***Figure 7F***). To validate the higher level of stress associated with sevoflurane GA, we conducted a series of tests to measure stress levels, including the open field test (OFT) (***Figure 7G***) and the elevated plus-maze test (EPM) (***Figure 7H***). Combining the results of the tests confirmed that the post-anesthesia period is associated with a high level of stress. To further determine the influence of chemogenetic inhibition of PVH^CRH^ neurons on the post-anesthesia stress response, we also performed stress-related behavioral testing on CRH-Cre mice expressing hM4Di receptors (n = 8 mice) in the PVH 30 min after i.p. administration of 3 mg/kg CNO or vehicle (***Figure 7A***). We observed that after 30 min of 1.6% sevoflurane exposure, mice moved shorter distances in the central area during the OFT (***Figure 7G and I***) and spent less time in the open arms during the EPM test (***Figure 7H and I***) compared to the pure oxygen inhalation group. Intriguingly, PVH^CRH^ neuron-inhibited mice showed similar central distance and time spent in the open arms to those of the pure oxygen inhalation group, indicating that chemogenetic inhibition of PVH^CRH^ neurons abolished the stress response after sevoflurane GA. To further explore whether mice developed post-operative delirium induced by sevoflurane which is characterized by disturbances in attention and cognition ^39,40^, if and how PVH^CRH^ neurons are involved in, we conducted the Y-maze test and novel object recognition test to test the cognitive function. We found that chemogenetic inhibition of PVH^CRH^ neurons ameliorated the short-term memory impairment caused by 30-minute exposure to sevoflurane GA (***Figure 7-figure supplement 9***), suggesting PVH^CRH^ neurons may involve in modulating sevoflurane-induced postoperative delirium.

## 3. Discussion

Here, applying cutting-edge techniques, we found that PVH^CRH^ neuronal activity decreased during sevoflurane induction and gradually increased during the emergence phase, followed by hyperactivities with increased stress levels in mice after termination of sevoflurane administration. Furthermore, the modulation of PVH^CRH^ neurons bidirectionally altered the induction and recovery of sevoflurane GA. More importantly, chemogenetic inhibition of this population attenuated the post-anesthesia stress response, supporting the role of PVH^CRH^ neurons in mediating states of consciousness and stress response associated with sevoflurane GA.

The reversible loss of consciousness induced by sevoflurane GA has been reported to be the result of mutual regulation of numerous nuclei, including several wake-promoting ensembles^41^, such as dopaminergic neurons in the ventral tegmental area ^42–45^, glutamatergic neurons in the PVT ^46^ and PB ^10,47^, and neurons expressing dopamine D1 receptors (D1R) in the nucleus accumbens (NAc)^36^, as well as sleep-promoting populations, such as GABAergic neurons in the rostromedial tegmental nucleus ^48,49^. In our study, we identified a new brain region involved in sevoflurane GA - PVH^CRH^ neurons - the activation of which prolongs the induction time and shortens the emergence time of sevoflurane GA, and modulates the neural inertia of the central nervous system (a tendency to resist behavioral state transitions between conscious and unconscious states)^50^. EEG is used to monitor the depth of anesthesia, and the EEG activity is linked to behavioral states of GA ^51^. Burst suppression is a phenomenon that occurs at deep anesthesia and is caused by the interaction of the hyperexcitable cortex with a refractory phase after the burst ^52,53^. This hyperexcitability is favored by reduced inhibition of anesthetics ^52^. Notably, photostimulation of PVH^CRH^ neurons effectively elicited cortical activation and behavioral emergence during steady sevoflurane GA state but only caused cortical activation during burst-suppression oscillations. Compared to the potent effect of optical stimulation of NAc D1R neurons during burst-suppression ^36^, our results indicate that optogenetic activation of PVH^CRH^ neurons is insufficient to reverse the reduced inhibition. Furthermore, similar to the ‘Arousal-Action Model,’ ^54^ GA-inhibiting neurons can also be classified into two categories referred to as ‘arousal neurons’ and ‘action neurons.’ Activation of the arousal neurons promotes EEG desynchronization but is insufficient to cause EMG activation during GA. In contrast, stimulation of action neurons causes both EEG desynchronization and behavioral emergence during GA. Based on this model, PVH^CRH^ neurons can be classified as the arousal neurons, while NAc D1R neurons belong to the action neurons. Furthermore, it has been reported that preoperative anxiety increases the anesthetic requirements intraoperatively and prolongs recovery from anesthesia ^55,56^, indicating that stress may induce abnormal neuronal activity, which influences the effect of anesthetic drugs and alters the induction of and emergence from GA. Given the vital role of PVH^CRH^ neurons in the stress response regulation, it is also possible that PVH^CRH^ neurons modulate sevoflurane GA through influencing stress levels.

The postoperative stress response is commonly associated with GA combined with surgery trauma. HPA activation during and after surgeries can result from multiple factors aside from anesthesia, including pain, anxiety, inflammation. Effective modulation of the stress response is of great importance in reducing the incidence of complications. As a predominant presentation of the post-anesthesia stress response, sevoflurane-induced emotional agitation is common, with an incidence of up to 80% in clinical reports, and is characterized by hyperactivity, confusion, and delirium ^57,58^. Such responses pose the risk of neurotoxicity, delayed neurological development, and even cognitive deficits. However, a broad consensus on animal models mimicking this post-anesthesia stress response has yet to be established. Stress often leads to grooming and other repetitive behaviors, such as digging and circling in rodents ^30,59,60^, and self-grooming is frequently observed and believed to contribute to post-stress de-arousal with adaptive value ^31,32^. In this study, we observed increased self-grooming and a series of other stress-related behavioral parameters during the post-anesthesia period, combined with an elevated serum CORT level, strongly indicating the high stress level, and supporting these measures as indicators of post-anesthesia stress responses in animal models. Of note, the hyperactivity of PVH^CRH^ neurons and behavior (e.g., excessive self-grooming and impaired cognitive function) in mice could partly recapitulate the observed agitation and delirium during the emergence from sevoflurane GA in patients. Our findings may further help us to understand the underlying mechanism related to post-anesthesia stress responses observed in patients, leading to the development of strategies to prevent or treat the post-anesthesia stress response.

Of particular note, the PVH is canonically viewed as the key endocrine controller of the stress response ^61^ and complex behavior after stress ^62,63^, including self-grooming ^64^. A recent study found that this role of the PVH was instructed by glutamatergic inputs from the lateral hypothalamus (LH) ^65^. Moreover, multiple lines of evidence have also demonstrated that PVH^CRH^ neurons are not only responsible for launching the endocrine component of the mammalian stress response ^61^ but also for orchestrating post-stress behavior, including grooming ^63^. For example, empirical evidence proposed that the ‘‘PVH^CRH^-the caudal part of the spinal trigeminal nucleus-spinal’’ descending pathway translates stress-related stimuli into motor actions of self-grooming ^66^. In our study, we observed elevated PVH^CRH^ neuronal activity along with excessive self-grooming during the post-anesthesia period. Inspiringly, chemogenetic inhibition of PVH^CRH^ neurons significantly alleviated these responses, underscoring the crucial role of PVH^CRH^ neurons in sevoflurane GA-induced self-grooming. Across the whole GA process, PVH^CRH^ neurons were greatly suppressed under sevoflurane GA but were recruited to control stress responses with rebound excitation after cessation of sevoflurane GA, and therefore evoked self-grooming and other behaviors to facilitate the reduction in stress level. Therefore, our data contribute to an emerging view that PVH^CRH^ neurons also play a key role in mediating the post-anesthesia stress responses.

Interestingly, we found that sevoflurane GA-induced self-grooming exhibited exclusive features different from other grooming models, and the LH, which provides glutamatergic inputs to the PVH in grooming ^65^, was not activated during the post-anesthesia period (***Figure 1-figure supplement 2B***), implying the existence of other functional pathways. PVH^CRH^ neurons received direct long-range GABAergic inputs from multiple brain regions, such as the LS, raphe magnus nucleus, and bed nucleus of the stria terminalis ^67^. Meanwhile, it has been demonstrated that the neuronal activity of the suprachiasmatic nucleus negatively regulates the activity of PVH^CRH^ neurons by GABA release ^68^. This evidence provides a possible explanation for the hyperactivity of PVH^CRH^ neurons in the post-anesthesia period, whereby sevoflurane GA suppresses activity of these brain areas, and therefore disinhibits PVH^CRH^ neurons by decreasing GABAergic inputs. Further research is required to fully understand this mechanism.

In conclusion, we have identified a significant node modulating the effect of sevoflurane general anesthesia in both during- and post-anesthesia stages: the PVH^CRH^ neurons. These neurons play a potent role in modulating anesthesia states in sevoflurane GA and are part of sevoflurane’s anesthesia regulatory network.

## 4. Experimental Section

### Mice

CRH-Cre mice (Crh < Cre > B6(Cg)-Crh^tm1(cre)Zjh^/J, JAX #012704) mice were kindly provided by Prof. Gang Shu from South China Agricultural University and were obtained from the Shanghai Model Organisms Center. Wild-type C57BL/6 and CD-1 mice were obtained from the Experimental Animal Management Department, Institute of Family Planning Science, Shanghai. Mice were group-housed in a soundproof room, with an ambient temperature of 22 ± 0.5°C and a relative humidity of 55% ± 5%. The mice were provided with adequate amounts of food and water, and were housed under an automatic 12-h light/12-h dark cycle (illumination intensity »100 lux, lights on at 07:00). Adult C57BL/6 male mice (CRH-Cre and wild-type mice, 8–10 weeks old, 20–30 g) were used for all experiments; adult CD-1 male mice (4–6 months old, 40–50 g) were used for the direct attack-induced model. All behavioral experiments were performed during the daytime (7:00–19:00). Animals were randomly divided into the experimental groups and control group. CRH-Cre mice were used in calcium fiber photometry recordings, chemogenetic and optogenetic manipulations, and wild-type mice were used for the identification of active neurons and the characterization of self-grooming during the post-anesthesia period. All experimental procedures were approved by the Animal Care and Use Committee of Fudan University (certificate number: 20160225-073). All efforts were made to minimize animal suffering and discomfort and to reduce the number of animals required to produce reliable scientific data.

### Surgery

Mice were anesthetized with 2% pentobarbital sodium (45 mg/kg) by intraperitoneal injection and placed on a stereotaxic frame (RWD Life Science, China). The temperature of each mouse was maintained constant using a heading pad during the operation. After shaving the head and sterilizing with 75% ethanol, the parietal skin was cut along the sagittal suture and small craniotomy holes were made above the PVH. For chemogenetic experiment, the virus containing recombinant AAV-EF1α-DIO-hM3D(Gq)-mCherry-WPRE (3.3 × 10^12^ vector genomes (VG)/mL, Brain VTA, China), or AAV2/9-hEF1α-DIO-hM4D(Gi)-mCherry-pA (3.3 × 10^12^ VG/mL, Taitool Bioscience, China), or a viral vector encoding Cre-dependent expression of Caspase-3 (AAV-CAG-FLEX-taCasp3-TEVp, 5.88 × 10^12^ VG/mL) was bilaterally injected into the PVH (anteroposterior [AP] = –0.4–0.5 mm, mediolateral [ML] = ± 0.2 mm, dorsoventral [DV] = 4.5 mm deep relative to endocranium) via a glass injection needle and an air-pressure-injector system at a rate of 0.1 μL/min. Control mice received injections of the control virus AAV-EF1α-DIO-mCherry-WPRE-hGH pA (5.35× 10^12^ VG/mL, Brain VTA, China) at the same coordinates and volumes. After 7 min of injection to each site, the glass pipette was left at the injection site for 8 min before slowly withdrawing.

For *in vivo* fiber photometry recordings and optogenetic experiments, after injecting the virus (AAV2/9-hSyn-DIO-jGCaMP7b-WPRE-pA, AAV2/9-hEF1a-DIO-hChR2-mCherry-pA, or AAV2/9-hEF1a-DIO-mCherry-pA, 1.61 × 10^13^ VG/mL, Taitool Bioscience, China) into the bilateral PVH at a rate of 0.1 μL/min for 7 min, the optical fiber cannula (outer diameter [OD] = 125 mm, inner diameter = 400 μm, numerical aperture [NA] = 0.37; Newdoon, Shanghai, China) was placed above the unilateral PVH ([AP] = –0.4–0.5 mm, [ML] = + 0.2 mm, [DV] = 4.0 mm deep relative to the endocranium) before implanting the EEG and EMG electrodes. The cannulas and the EEG and EMG electrodes were secured to the skull with dental cement. Mice were then kept in a warm environment until they resumed normal activity and fed carefully for 2 weeks before the following experiments.

### Calcium fiber photometry recordings

Using a multichannel fiber photometry system equipped with 488-nm laser (OBIS 488LS; Coherent) and a dichroic mirror (MD498; Thorlabs), the fluorescence signals of the GCaMP were collected by a photomultiplier tube (R3896, Hamamatsu) and recorded using a Power1401 digitizer and Spike2 software (CED, Cambridge, UK) with simultaneous EEG/EMG recordings. During this experiment, mice were first placed in a cylindrical acrylic glass chamber connected to a sevoflurane vaporizer (VP 300; Beijing Aeonmed, China). The concentration of sevoflurane was monitored by an anesthesia monitor (Vamos, Drager, China) connected to the chamber. In the dose-dependent experiments, after recording the baseline signals of awake mice during a 20-min adaption period, different concentrations of sevoflurane (1.2%, 1.6%, or 2.0%) were continuously delivered by 1.5 L/min flow of 100% oxygen for 20 min. The group inhaling 100% oxygen was set as the control group. As for the observation of the Ca^2+^ signals during the post-anesthesia period, the baseline signals of awake mice were recorded for 30 min before 30-min delivery of 1.6% of sevoflurane. After turning off the sevoflurane vaporizer, the recording lasted until the Ca^2+^ signals returned to the baseline level.

### Chemogenetic manipulations

All the chemogenetic experiments were performed between 19:00–24:00, with at least a 24-h washout time. After intraperitoneally pretreating the animals with saline or 3 mg/kg of CNO (C4759, LKT, USA), the mice were placed in a cylindrical acrylic glass chamber with 1.5 L/min flow of 100% oxygen for 20 min for habituation. To determine the sensitivity of sevoflurane, mice experienced stepwise increasing exposure to sevoflurane, initially delivered at 1.2% and increased every 10 min in increments of 0.1%. The righting reflex of each animal was assessed after 10 min at each concentration, ensuring both brain and chamber balance. If a mouse successfully returned to the prone position twice in a row, then it was considered to have a complete righting reflex. Otherwise, the righting reflex was determined not to be present. The induction and emergence experiments were conducted as follows. Each mouse was intraperitoneally administered saline or CNO (3 mg/kg, i.p.) 1 h before the experiment. Next, 30-min 2% sevoflurane was delivered with 1.5 L/min 100% oxygen after 20-min habituation. Following exposure to 2% sevoflurane for 30 min, the mouse was removed from the glass chamber and kept in the supine position in the indoor air. The time to induction was defined as the interval between the time when the mouse was put into the glass chamber full of 2% sevoflurane to the time when the mice exhibited LORR. The time to emergence was defined as the interval between the time when the mouse was removed from the chamber to the time when it regained the righting reflex with all four paws touching the floor. Throughout the entire experimental period, an electric blanket was put under the chamber to maintain the chamber temperature at 36°C.

### Optogenetic stimulation

Optical stimulation was first performed during induction of and recovery from sevoflurane GA. After recording an awake EEG/EMG baseline for 10 min (continuously inhaling pure oxygen), we increased the concentration of sevoflurane in the chamber to 2.0% and applied light-pulse trains (5-ms pulses at 30 Hz) simultaneously until LORR. Next, after each mouse inhaled 2.0% sevoflurane for 30 min, we turned off the sevoflurane vapor and initiated acute light stimulation (5-ms pulses at 30 Hz) until RORR.

For manipulation during the steady sevoflurane GA state, light-pulse trains (473-nm, 5-ms pulses at 30 Hz for 60 s) were applied according to a previous protocol ^44^. In brief, a baseline EEG/EMG was recorded after habituation in the acrylic glass chamber for 10 min. Next, the sevoflurane vapor was turned on and the mice were subjected to sevoflurane with an initial 2.5% sevoflurane for 20min and then placed in a supine position and then the sevoflurane concentration was reduced to 1.4%. If the animals showed any signs of LORR, the concentration was increased by 0.1% until the LORR maintained at a constant concentration for at least 20 min. . Photostimulation (5-ms pulses at 30 Hz for 60 s) was applied through a laser stimulator (SEN-7103, Nihon Kohden, Japan) when EEG patterns showed characteristics of steady sevoflurane GA state (low-frequency, high-amplitude activity) for at least 5 min. As for the burst-suppression period, a baseline EEG/EMG was recorded during habituation in the acrylic glass chamber for 10 min. Next, we delivered 2% sevoflurane with pure oxygen at a speed of 1.5 L/min for induction initially. After entering a continuous and steady burst-suppression oscillation mode for at least 5 min, we applied blue-light stimulation (5-ms pulses at 30 Hz) for 60 s. A power meter (PM10, Coherent, USA) was used to measure the power intensity of the blue light before experiments.

### EEG/EMG analysis

SleepSign software (Kissei Comtec, Japan) was used to record and analyze all EEG/EMG signals. EEG/EMG signals were magnified and filtered (EEG, 0.5–30 Hz; EMG, 20–200 Hz), and were digitized at a 128 Hz sampling rate. Subsequently, fast-Fourier transformation was used to calculate the EEG power spectra for successive 4 s epochs in the frequency range of 0–25 Hz. EEG frequency bands (delta, 0.5–4.0 Hz; theta, 4.0–10 Hz; alpha, 10–15 Hz; beta, 15–25 Hz) were established in view of previous research, and relative variations in aggregate power were calculated. In the optogenetic experiments during steady sevoflurane GA state, a total of 120-s EEG signals were analyzed (before and during acute light stimulation). As for optogenetic tests during burst-suppression oscillations, a total of EEG signals from 180-s epochs were calculated (before, during, and after acute light stimulation). The BSR was calculated by the percentage of suppression in the time spans of 1 minute before, during, or after photostimulation. Raw EEG data recorded by SleepSign software were converted to text format for further analysis of BSR using MATLAB as demonstrated previously^36^. The minimum duration of burst and suppression periods was set to 0.5 s.

### Immunohistochemistry

The immunohistochemistry test was conducted according to the previous studies ^69^. To confirm the correct injection sites, after finishing all related experiments, each mouse was anesthetized with chloral hydrate (360 mg/kg) and then perfused intracardially with 30 mL phosphate-buffered saline (PBS) followed by 30 mL 4% paraformaldehyde (PFA). The brains of the mice were removed and postfixed in 4% PFA overnight and then incubated in 30% sucrose phosphate buffer at 4°C until they sank. Coronal sections (30 μm) were cut on a freezing microtome (CM1950, Leica, Germany), and the fluorescence of injection sites was examined to locate the PVH according to the histology atlas of Paxinos and Franklin (2001, The Mouse Brain in Stereotaxic Coordinates 2nd edn [San Diego, CA: Academic]) using a microscope (Fluoview 1200, Olympus, Japan).

For immunostaining of c-fos, mice were sacrificed at set time according to different experiments and perfused with PBS followed by 4% PFA in 0.1-M phosphate buffer. The brain was then dissected and fixed in 4% PFA at 4°C overnight. Fixed samples were sectioned into 30-μm coronal sections using a freezing microtome (CM1950, Leica, Germany). Brain sections were washed in PBS three times (5 min each time) and incubated for 48 h with primary antibody in PBST (containing 0.3% Triton X-100) on a 4°C agitator using rabbit anti-c-fos (1:8000, Abcam, USA) antibody, or rabbit anti-CRF (1:800, Immunostar 20084, bilateral intracerebroventricular infusion of colchicine (total 20 μg in 500 nL saline, coordinates from Bregma: AP = -0.22 mm, ML = ± 1.15 mm, DV = 2.06 mm) 24 h before perfusion)^70^. Next, brain sections were washed in PBS three times (5 min each time) and incubated with a donkey-anti-rabbit Alexa488 secondary antibody (1:1000, Abcam, USA) in PBST for 2 h at room temperature. Finally, the sections were mounted on glass slides, dried, dehydrated, and cover-slipped. Fluorescence images were collected with an Olympus VS120 automated slide scanner. Only data from mice in whom the AAV infection and optical fiber location were confirmed were included. The number of animals that were excluded because of poor virus expression or misplaced optical fibers are as follow: jGCaMP7b, n = 5; hM3Dq, n = 7; hM4Di, n = 10; mCherry, n = 5; ChR2, n = 12; wrong optical fiber location, n = 12.

### Behavior experiments Righting reflex assessment

GA was defined by the loss of the righting reflex, a widely used surrogate measure in rodents. All mice were placed in the air-tight, temperature-controlled, 200 mL cylindrical chambers with 100% oxygen flowing at a speed of 1.5 L/min for 10 min for the sake of habitation. To determine the EC_50_ of LORR, the initial concentration of sevoflurane was set at 1.0%, and the concentrations were increased by 0.1% every 15min. During the last 2 min of every 15 min, the chambers were rotated 180° to observe whether the righting reflex of the mice was present, and the number of animals that lost their righting reflex at each concentration was recorded. To determine the EC_50_ of RORR, all the mice were anesthetized with 2.0% sevoflurane for 30 min, and the concentration was decreased in decrements of 0.1% until all the mice had recovered their righting reflex. The number of mice that had recovered their righting reflex at each concentration was counted. Mice experienced stepwise increasing and diminishing exposure to sevoflurane, with 15 min taken at each step, during which and an electric blanket was used to maintain the chamber temperature at 36°C. The righting reflex of each animal was assessed after 15 min at each concentration, ensuring both brain and chamber balance. If a mouse successfully returned to the prone position twice in a row, then it was considered to have a complete righting reflex. Otherwise, the righting reflex was determined not to be present.

### Arousal Scoring

Arousal responses were scored according to the methods of previous studies^36^. Spontaneous movements of the limbs, head, and tail were scored by the movements’ intensity as three levels: absent (0), mild (1), or moderate (2). Righting was scored as 0 if the mouse remained LORR, and 2 if the mouse recovered its righting reflex. Walking was scored as follows: 0 = no further movements; 1 = crawled without raising the abdomen off the chamber bottom; and 2 = walked with all four paws and the abdomen off the chamber bottom. The total score for each mouse was defined as the sum of all categories during the photostimulation period.

### Assessment of self-grooming

The behavior of the animals was recorded in a test chamber (cylindrical acrylic glass, 15-cm diameter, 20-cm height) in a quiet and sound-proof room by a video camera (HIKVISION, US). Mice were habituated in the test chamber for 20 min before testing. For the spontaneous grooming model, the mouse was placed into the test chamber and spontaneous activities were recorded over a 20-min period. For restraint stress-induced grooming, the mouse was restrained in a stainless-steel tube (5-cm diameter, 10 cm length) for 20 min and then placed immediately into the test chamber for 20-min video recording. For direct attack-induced grooming, a C57BL/6 mouse was placed directly into the home cage of a larger and aggressive CD-1 mouse for 5 min with the aggressor present ^71^. Immediately afterward, the C57BL/6 mouse was placed in the test chamber, and its behavior was recorded for 20 min. For water spray-induced grooming ^65^, the mouse was sprayed with a spray bottle prefilled with sterile water (25°C) directed to the face, belly, and back, before placing the mouse in the test chamber for 20 min of video recording. For swimming-induced grooming, the mouse was placed into a plastic cylinder filled with water at 22–26°C and allowed them to freely swim for 5 min. Excess water was then removed before the animal was placed into the test chamber for video recording for 20 min.

The self-grooming behavior was manually quantified by observers blinded to the experimental conditions. Self-grooming was defined as when the mouse licked its own body parts, including the paws legs, tail, and genitals, or stroked its own nose, eyes, head, body, with the front paws ^32^. An interruption of 6 s or more separated two individual bouts ^34,35^. The number of grooming bouts, the mean bout duration, and the total grooming time spent in the test period were evaluated. For the analysis of the patterns and microstructures of grooming behavior, the time spent on grooming different body parts and the transitions between different phases were calculated based on previous protocols ^34,35,72^. The following scaling systems based on grooming stages were used: no grooming (0), paw licking (1), nose/face/head wash (2), body grooming (3), leg licking (4), and tail/genitals grooming (5). Correct transitions were defined as the following progressive transitions between the grooming stages: 0–1, 1–2, 2–3, 3–4, 4–5, and 5–0. Incorrect transitions were characterized by skipped (e.g., 0–5, 1–5) or reversed (e.g., 3–2, 4–1, 5–2) stages. Interrupted grooming bout was considered if at least one interruption was recorded during the transition.

### Open field test

The OFT is one of the most widely used behavioral tests to evaluate anxiety-like behavior in rodents ^72^. In the OFT, a camera was used to record the movement of mice in a 50 x 50-cm area. The middle square (30 x 30 cm) of the box was considered the central area. The entire device was surrounded by 40-cm high walls and its inner surface was painted black. In a quiet and sound-proof room with dim light (≈ 20 lx), a mouse was placed in the middle of the arena and allowed to move freely for 10 min. An automatic video tracking system (Tracking Master V3.0, Shanghai Vanbi Intelligent Technology Co., Ltd.) was used to record the ambulatory activity of the mice ^73^. After each test, the apparatus was wiped clean with 75% alcohol to avoid the influence of the previous mouse. The total and central travel distances were quantified.

### Elevated plus-maze test

The EPM test is commonly used to measure anxiety-like behavior in mice ^72^. The device consists of two open arms (20 x 6 x 0.5 cm) and two closed arms (20 x 6 x 30 cm), which lie across each other with an intersection part of 6 x 6 x 0.5 cm. The open arms have insignificant walls (0.5 cm) to reduce the frequency of falls, whereas the closed arms are surrounded by high walls (30 cm). The whole device was 70-cm above the floor. The camera was placed above the EPM in the behavioral test room. Each mouse was placed in the behavioral testing room 1 h before the test for adaption. During the test, the mouse was placed in the intersection area of the maze, facing an open arm. After exploring the maze freely for 5 min, the mouse was removed from the EPM and returned to its home cage. The maze was cleaned with a spray bottle and paper towel before testing the next mouse. An automatic video tracking system (Tracking Master V3.0, Shanghai Vanbi Intelligent Technology Co., Ltd.) was used to analyze the movement data of the mouse. An entry was defined when half of the mouse’s body entered an area.

### Y-maze test

The Y-maze test for evaluating learned behavior was conducted as previously described^39,40^. Initially, the test mouse was placed at the start of one arm, facing away from the center, and allowed to explore the apparatus freely for 5 minutes while another arm was blocked. After this period, the mouse was returned to its home cage. Subsequently, the block was removed, and the mouse was allowed to explore all arms freely for another 5 minutes. The discrimination index, which is the time spent in the novel arm divided by the time spent in both arms, was used to assess spatial memory.

### Novel object recognition test

The novel object recognition test was conducted in the same arena as the open field test (OFT). During the habituation session, the test mouse was placed in the arena to explore for 10 minutes. Following habituation, two objects of similar size but different shape and color were placed in opposite corners of the arena. The test mouse was then placed in the center and allowed to explore for 10 minutes. After this exploration, all mice were exposed to 2.0% sevoflurane for 30 minutes and then returned to their home cages. Thirty minutes later, saline or CNO was injected. One hour after the injection, one of the objects was replaced, and the same test mouse was allowed to explore for another 10 minutes. Interactions with each object (defined as sniffing and/or having the head within 2-3 cm of the object) were recorded for analysis. Mice that did not reach a minimum of 20 seconds of exploration time for both objects during either of the two 10-minute trials were excluded from analysis, as it could not be confirmed that they spent enough time exploring to learn or discriminate between the objects. The time spent interacting with the novel object, divided by the time spent interacting with both objects, was used to describe the preference for the novel object.

### Measurement of CRH and CORT levels

Mice were decapitated immediately at one hour after the mice received either saline or CNO (3 mg/kg, C2041, LKT) treatment, respectively. Blood was rapidly collected from the retroorbital plexus of the mice subjected to sevoflurane GA or pure oxygen. Considering the circadian rhythm release pattern of CORT, which has lower concentrations in the morning ^74^, blood sampling was conducted between 8:00 and 12:00 to minimize the effect of time of day on glucocorticoid levels. Blood samples were allowed to clot for 2 h at room temperature before centrifugation for 15 min at 1000 ×g. Serum was removed and assayed immediately or aliquoted, and samples were stored at −80°C. The concentrations of CRH and CORT in the serum were detected using enzyme-linked immunosorbent assay (ELISA) kits (CSB-E14068m and CSB-E07969m, CUSABIO Technology, China) according to the manufacturer’s instruction.

### Quantification and statistical analysis

All data are presented as the mean ± SD. The selection of sample sizes was based on similar experiments that use chemogenetic and optogenetic methods ^10,36^. Paired or unpaired two-tailed Student’s t-test was selected to compare the two groups. If the data did not satisfy the normal distribution, further analysis was performed using Wilcoxon signed-rank tests or Mann–Whitney rank-sum tests. Multiple comparisons were made using one-way repeated-measures ANOVA with Tukey’s post-hoc test or two-way repeated-measures ANOVA followed by Sidak’s post hoc comparison test. Statistical analysis was performed using GraphPad Prism 8.0 (GraphPad Software, USA). Full details of the statistical analyses are shown in the figure legends. *p* < 0.05 was statistically significant in all cases.

## Acknowledgements

We thank Prof. Gang Shu from South China Agricultural University for kindly providing CRH-Cre mice, and Yuan Meng, Mingxuan Yang, Ziheng Zhao, Yingfan Guo and Yi Zhang for assisting experiments and data analysis.

## Author contributions

Shan Jiang and Lu Chen contributed equally to this work. Shan Jiang conceived, designed, and performed the experiments, as well as analyzed/interpreted the data, prepared the figures, and wrote the manuscript. Lu Chen performed the experiments, analyzed/interpreted the data and prepared the figures. Chang-Rui Chen and Zhi-Li Huang designed the experiments and revised the manuscript. Chang-Rui Chen, Wei-Min Qu and Zhi-Li Huang supervised the study and acquired funding.

## Data availability statement

The data that support the findings of this study are available from the corresponding author upon reasonable request.

