## Supplementary File for "Hypothalamic CRH Neurons Modulate Sevoflurane Anesthesia and The Post-anesthesia Stress Responses"

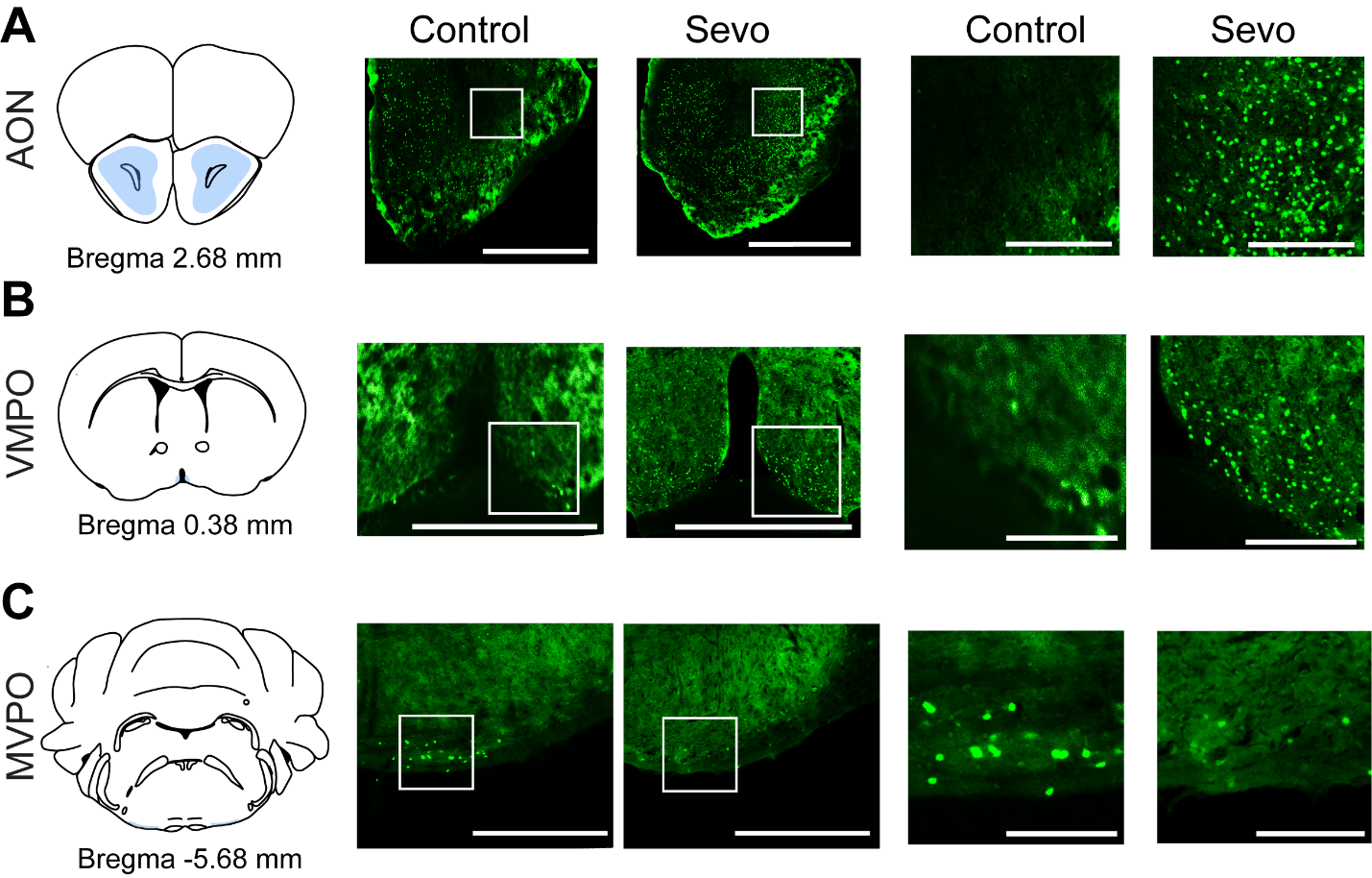

Figure 1-figure supplement 1. Whole-brain mapping of active neurons during the post-anesthesia period.

**A-C.** Representative images of c-fos staining in the AON, VMPO, and VMPO for a control and an experimental animal exposed to sevoflurane GA. Scale bar, 500 µm (left), 250 µm (right, enlarged). AON, anterior olfactory nucleus; MVPO, medioventral periolivary nucleus; VMPO, ventromedial preoptic nucleus.

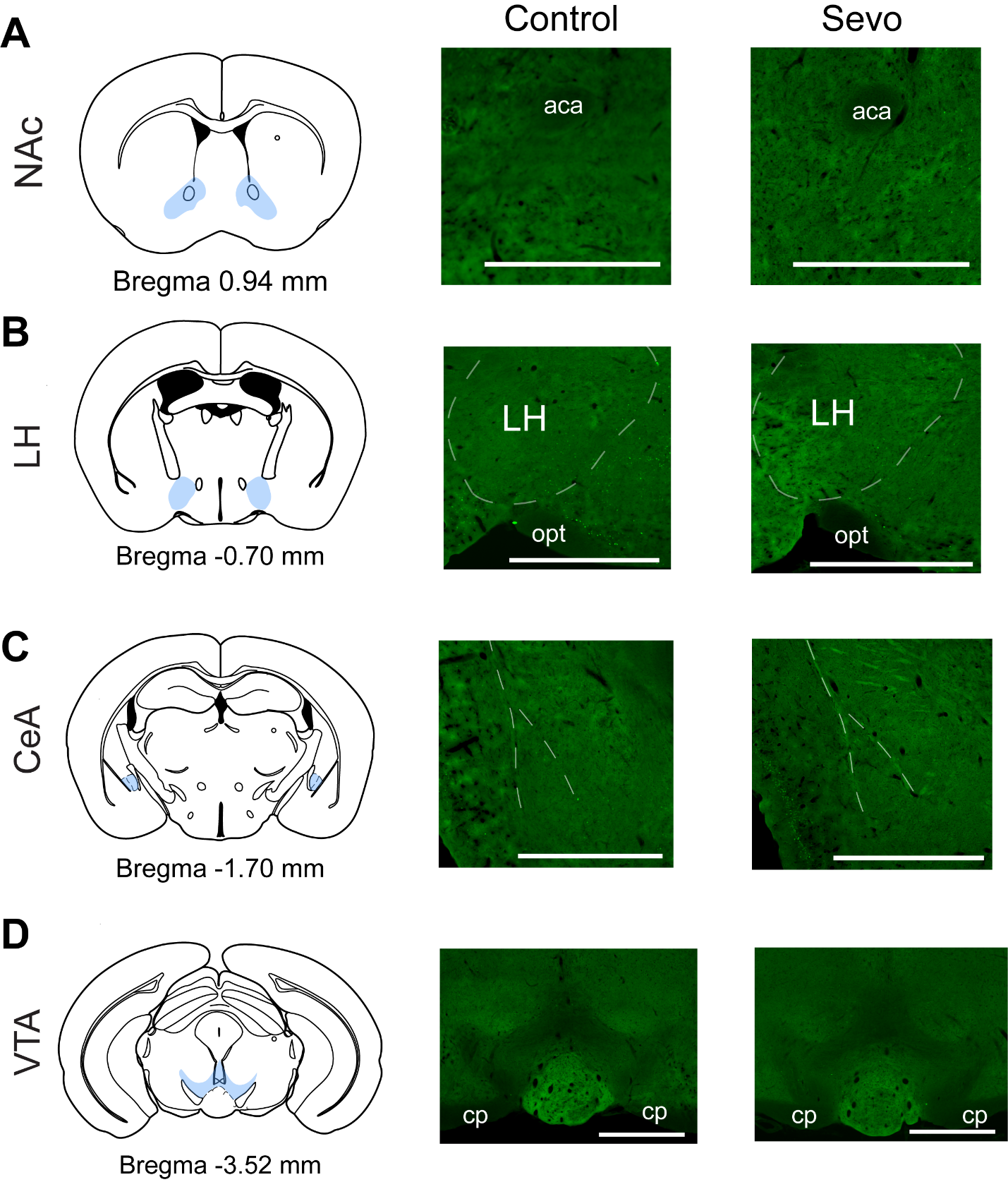

Figure 1-figure supplement 2. Representative images of brain regions without robust c-fos expression.

**A-D.** Representative images of brain regions without robust c-fos expression. aca, anterior commissure - the anterior part; CeA, central amygdaloid nucleus; cp, cerebral peduncle - the basal part; LH, lateral hypothalamus; NAc, nucleus accumbens; opt, optic tract; VTA, the ventral tegmental area. Scale bar, 500 µm.

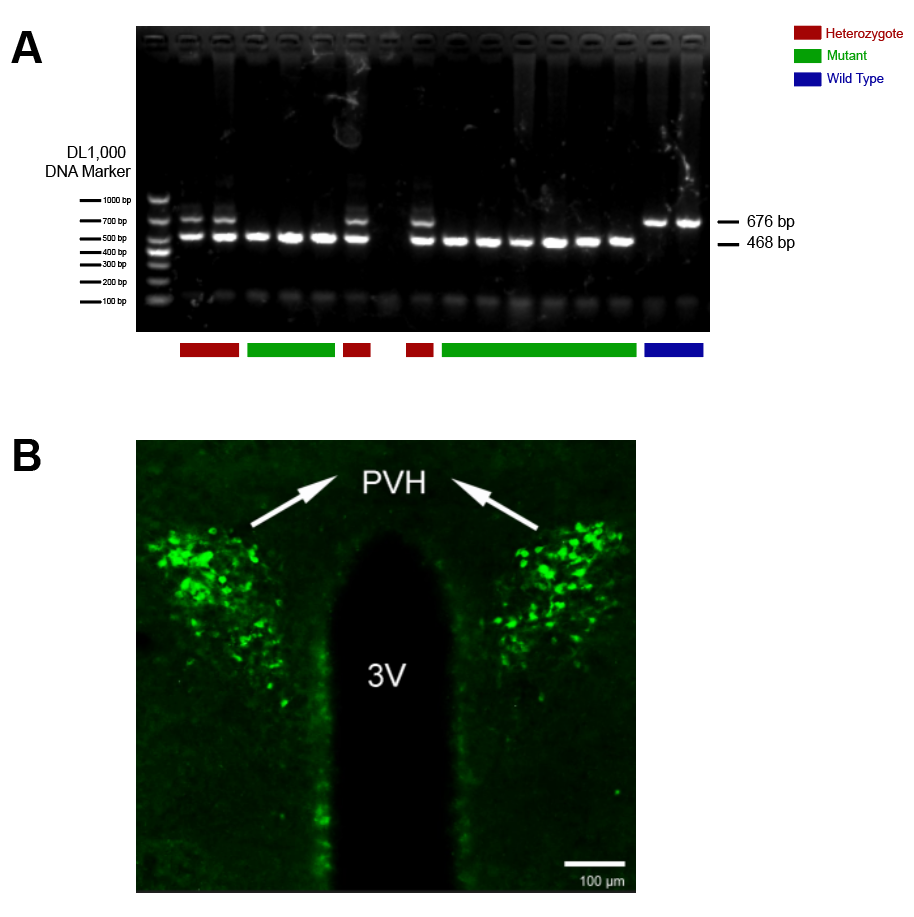

Figure 1- figure supplement 3. Validation of CRH-Cre mice and CRH antibody.

**A**. Standard PCR of CRH-Cre mice (B6(Cg)-Crhtm1(cre)Zjh/J). **B**. Representative images showing CRH immunoreactivity using CRH antibody at a concentration of 1:800, Scale bar = 100 μm. The 468-bp CRH-specific PCR product was amplified in mutant (CRH-Cre^+/+^) mice; in heterozygote (CRH-Cre^+/-^) mice, both the 468-bp and the 676-bp PCR products were detected; in wild type (WT) mice, only the 676-bp WT allele-specific PCR product was amplified.

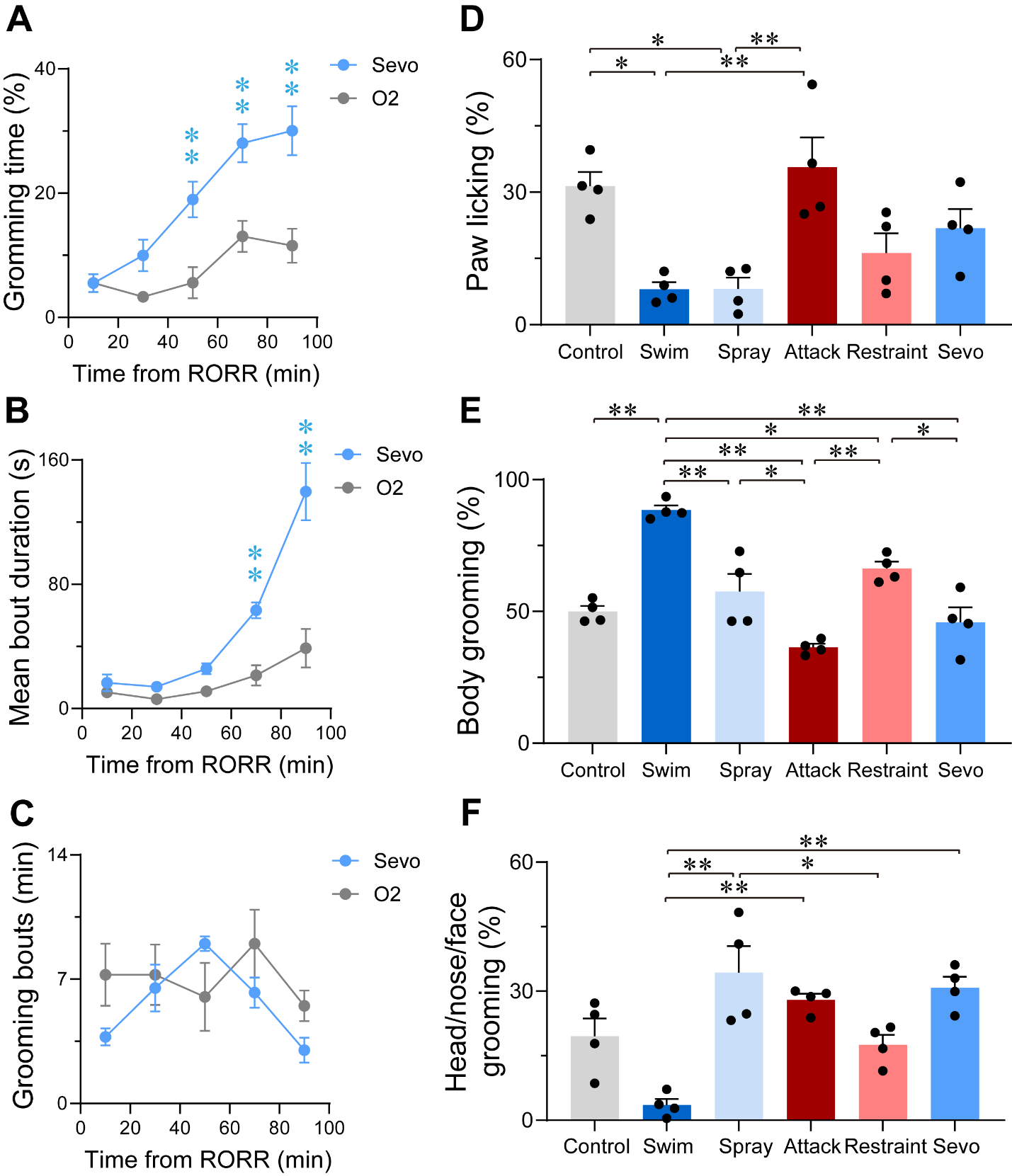

Figure 3-figure supplement 4. Characterization of sevoflurane GA-induced grooming.

**A-C.** Time courses for grooming time percentage (**A**, F (4, 24) = 4.268, *p* = 0.0095), mean bout duration (**B**, F (4, 24) = 17.42, *p* < 0.0001), grooming frequency (bouts per min, **C**, F (4, 24) = 1.927, *p* = 0.1385) within 100 min in mice after 30-min exposure of sevoflurane GA or oxygen (n = 4, two-way ANOVAs followed by Sidak’s test). **D-F.** Quantification of the number of bouts spent on paw licking (**D**, F (5, 18) = 7.824, *p* = 0.0005), body grooming (**E**, F (5, 18) = 21.67, *p* < 0.0001) and head/nose/face grooming (**F**, F (5, 18) = 10.72, *p* < 0.0001)) in the six grooming models (n = 4, one-way ANOVA with Tukey’s post-hoc test. **p* < 0.05; ***p* < 0.01).

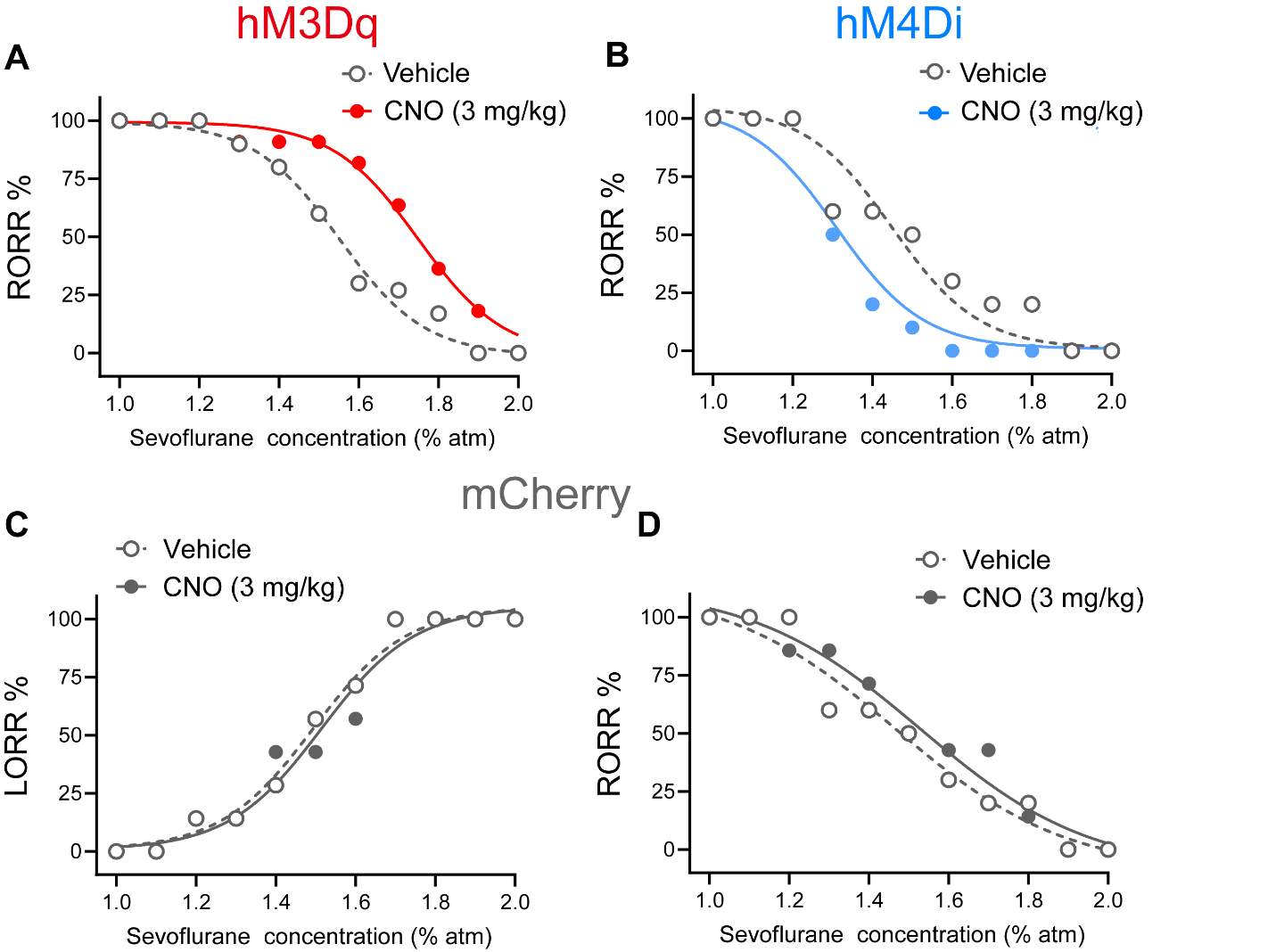

**Figure 4-figure supplement 5. Dose-response curves showing the percentages of mice exhibiting LORR (C) or RORR (A, B, D) in response to incremental or decreased sevoflurane concentrations for the PVH^CRH^ neurons activation (A), inhibition (B), and control (C-D) groups.** CNO, clozapine-N-oxide; LORR, loss of righting reflex.

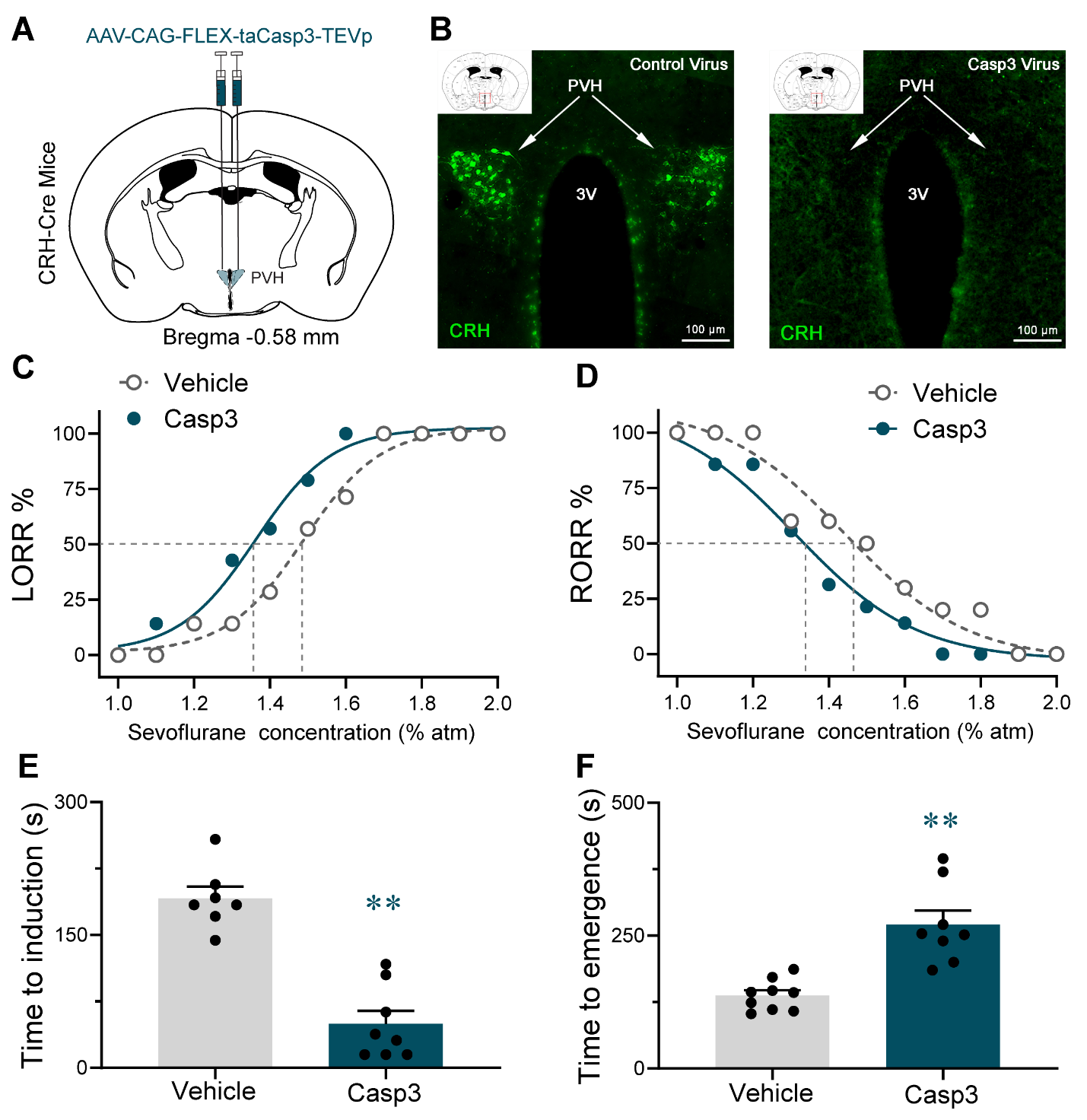
Figure 4-figure supplement 6. Lesion of PVH^CRH^ neurons facilitated induction of and delayed emergence from sevoflurane GA.

**A.** Schematic of AAV-CAG-FLEX-taCasp3-TEVp injected into the PVH of CRH-Cre mice. **B.** Representative coronal sections containing PVH regions from CRH- Cre mice sham-injected (Vehicle) or injected with AAV-CAG-FLEX-taCasp3-TEVp into the PVH. Scale bars, 100 μm. **C-D.** Dose-response curves showing the percentages of mice exhibiting LORR (**C**) or RORR(**D**) in response to incremental or decreasing sevoflurane concentrations for the vehicle and lesion groups. **E.** Induction time with 2% sevoflurane exposure of vehicle or lesion groups (n = 8, paired *t*-test, *p* < 0.0001). **F**. Emergence time with 2% sevoflurane exposure of vehicle or lesion groups (n = 8, paired *t*-test, *p* = 0.0002).

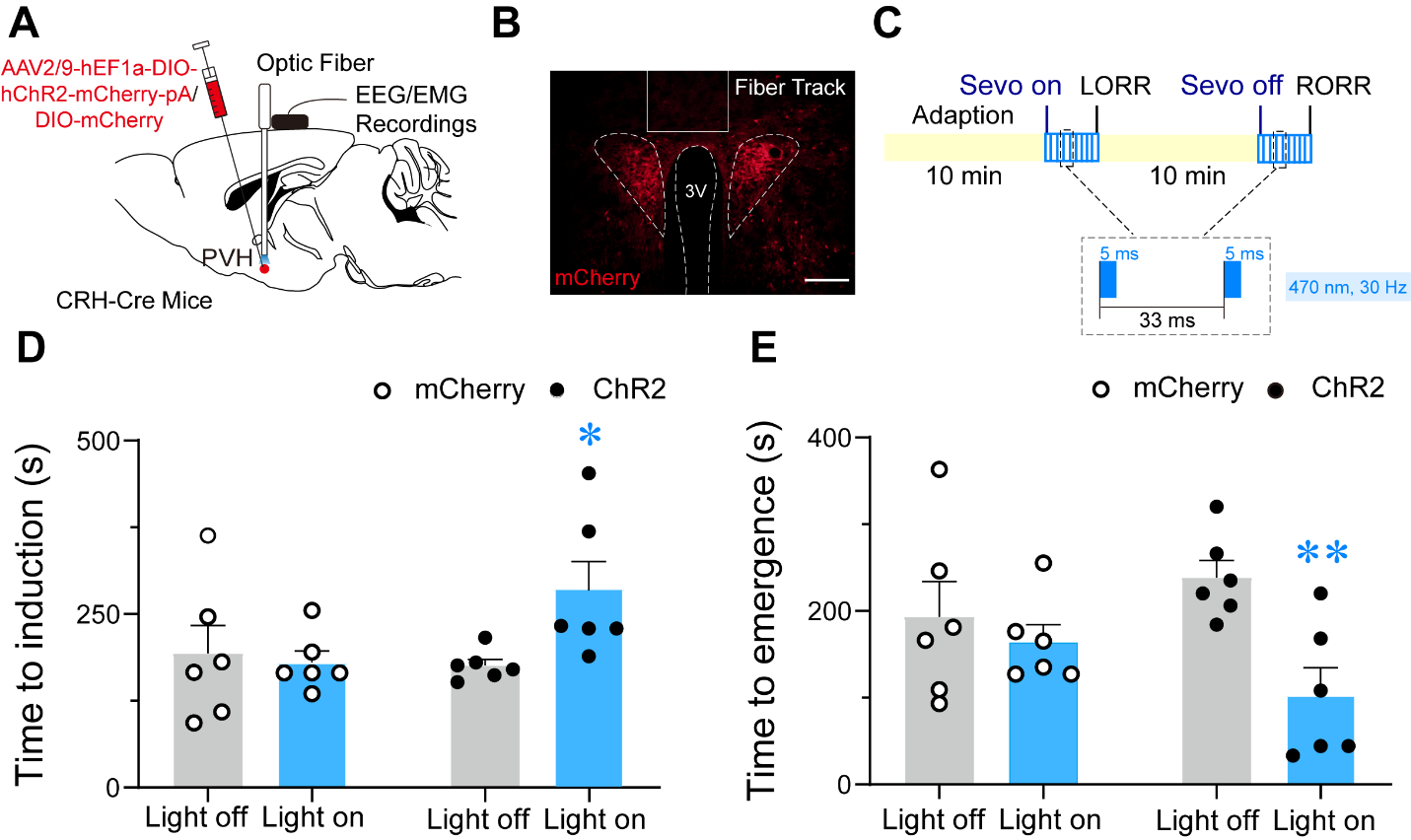

Figure 4-figure supplement 7. Optogenetic stimulation of PVH^CRH^ neurons delayed induction and facilitated emergence from sevoflurane GA.

**A.** Schematic of optogenetic stimulation of ChR2-expressing PVH^CRH^ neurons with EEG/EMG recordings. **B.** Image of ChR2-expressing PVH^CRH^ neurons (Bottom, scale bar: 200 μm). **C.** Protocol for optogenetic activation during sevoflurane GA. **D-E.** Optical activation of PVH^CRH^ neurons shortened the induction time (**D**) and prolonged the emergence time from 2% sevoflurane GA (**E**), mCherry-light-on vs ChR2-light-on, unpaired *t*-test; ChR2-light-on vs ChR2-light-off, paired *t*-test. n = 6, **p* < 0.05, ***p* < 0.01.

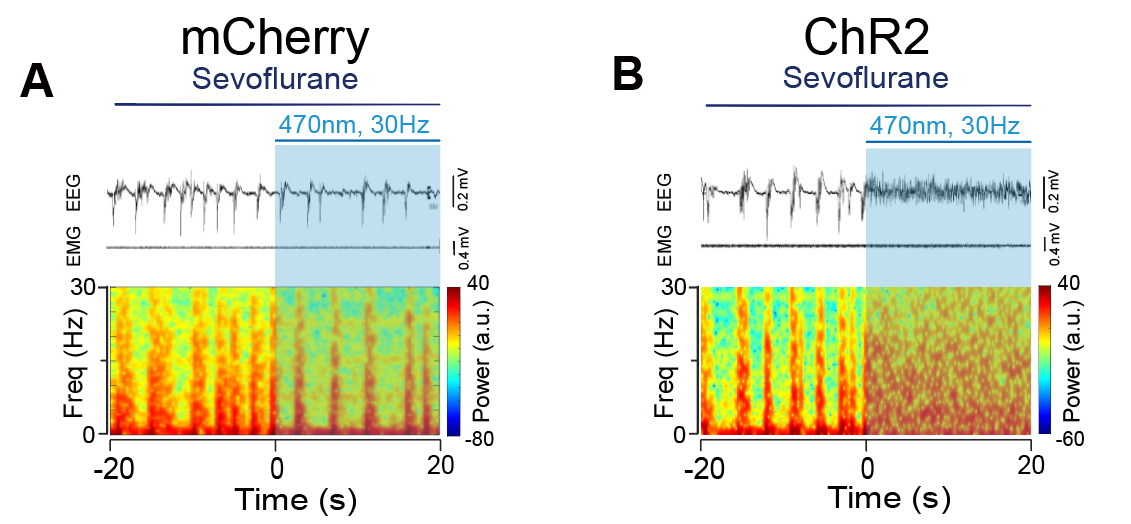

Figure 6-figure supplement 8. Optogenetic stimulation of PVH^CRH^ neurons induced cortical activation during burst-suppression oscillations induced by deep sevoflurane GA.

**A-B.**­ Typical examples of EEG, EMG, and EEG power spectral in a mouse injected with AAV-DIO-mCherry (**A**) or AAV-DIO-ChR2-mCherry (**B**) following acute photostimulation (30 Hz, 5 ms, 60 s) during burst-suppression oscillations. Time 0 indicates the beginning of photostimulation. The blue shadow indicates first 20 s of the 60-s duration of blue light stimulation.

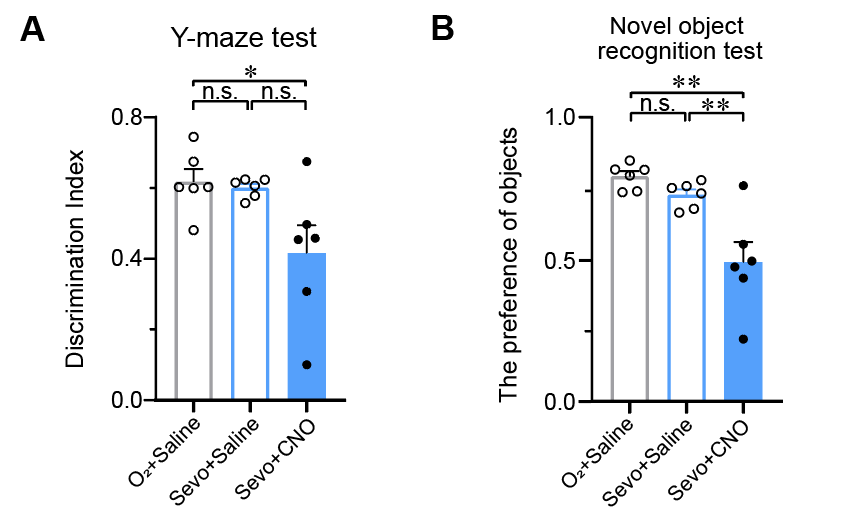

Figure 7-figure supplement 9. PVH^CRH^ neurons involved in modulating sevoflurane-induced short-term memory impairment.

**A-B**. Y-maze (**A**) and novel object recognition (**B**) tests after inhalation of sevoflurane or pure oxygen and administration of saline or CNO (**A**, n = 6, saline: O2 vs. sevo, *p* = 0.9700, t = 0.4477, df = 15, 95% CI = -0.1691 to 0.2028; O2 (Saline) vs. sevo (CNO), *p* = 0.0324, t = 2.325, df = 15, 95% CI = 0.01628 to 0.3882; sevo: saline vs. CNO, *p* = 0.0507, t = 2.318, df = 15, 95% CI = -0.0005627 to 0.3714; F (2, 15) = 4.914, *p* = 0.0228; (**B**, n = 6, saline: O2 vs. sevo, *p* = 0.5653, t = 2.462, df = 15, 95% CI = -0.09701 to 0.2262; O2 (Saline) vs. sevo (CNO), *p* = 0.0006, t = 4.105, df = 15, 95% CI = 0.1416 to 0.4648; sevo: saline vs. CNO, *p* = 0.0043, t = 3.227, df = 15, 95% CI = - 0.07702 to 0.4002; F (2, 15) = 13.18, *p* = 0.0005). Statistical comparisons were conducted using one-way ANOVA followed by Sidak’s tests. **p* < 0.05, ***p* < 0.01, n.s., no significant differences.

Figure 4-Table supplement 1. Mean value of EC50 for sevoflurane dose-response curves.

| EC_50_ (%) | mCherry | | Chemogenetic activation | | Chemogenetic inhibition | | Genetic ablation |
| --- | --- | --- | --- | --- | --- | --- | --- |
| LORR | | | | | | | |
| Mean | Vehicle | CNO | Vehicle | CNO | Vehicle | CNO |  |
| PVH^CRH^ neurons | 1.491 | 1.517 | 1.516 | 1.650 | 1.500 | 1.337 | 1.358 |
| 95% CI |  | | | | | | |
| PVH^CRH^ neurons | 1.444 to 1.536 | 1.292 to 1.690 | 1.509 to 1.522 | 1.616 to 1.682 | 1.496 to 1.504 | 1.297 to 1.368 | 1.292 to 1.406 |
| RORR | | | | | | | |
| Mean | Vehicle | CNO | Vehicle | CNO | Vehicle | CNO |  |
| PVH^CRH^ neurons | 1.420 | 1.571 | 1.538 | 1.791 | 1.424 | 1.306 | 1.326 |
| 95% CI |  | | | | | | |
| PVH^CRH^ neurons | 1.247 to 1.587 | 1.228 to 1.953 | 1.483 to 1.607 | 1.742 to 1.909 | 1.254 to 1.835 | 1.282 to 1.333 | 1.234 to 1.378 |

EC_50_: Alveolar concentration at which half of the mice lose their righting reflex, 95% CI: 95% confidence index.

Figure 5-Table supplement 2. Behavioral responses of CRH-Cre mice under sevoflurane steady-state GA during photostimulation.

| Group | Leg movement | Head movement | Tail  movement | Righting  reflex | Walking | Total  score |
| --- | --- | --- | --- | --- | --- | --- |
| 1-ChR2 | 2 | 2 | 2 | 2 | 2 | 10 |
| 2-ChR2 | 2 | 2 | 2 | 2 | 2 | 10 |
| 3-ChR2 | 2 | 2 | 1 | 2 | 2 | 9 |
| 4-ChR2 | 2 | 2 | 2 | 2 | 1 | 9 |
| 5-ChR2 | 2 | 2 | 2 | 2 | 0 | 8 |
| 6-ChR2 | 2 | 2 | 2 | 2 | 0 | 8 |
| 1-mCherry | 0 | 0 | 0 | 0 | 0 | 0 |
| 2-mCherry | 0 | 0 | 0 | 0 | 0 | 0 |
| 3-mCherry | 0 | 0 | 0 | 0 | 0 | 0 |
| 4-mCherry | 1 | 0 | 0 | 0 | 0 | 1 |
| 5-mCherry | 0 | 0 | 0 | 0 | 0 | 0 |
| 6-mCherry | 0 | 0 | 0 | 0 | 0 | 0 |

Leg, head, and tail movements, as well as states of righting reflex and walking in each mouse were determined during 60-s optical stimulation while mouse still inhaled with constant sevoflurane. The total score is the sum of the above categories.

Statistic report

**Figure 2E**

| Number of families | 1 |  |  |  |  |  |  |  |
| --- | --- | --- | --- | --- | --- | --- | --- | --- |
| Number of comparisons per family | 6 |  |  |  |  |  |  |  |
| Alpha | 0.05 |  |  |  |  |  |  |  |
| Tukey's multiple comparisons test | Mean Diff. | 95.00% CI of diff. | Significant? | Summary | Adjusted P Value |  |  |  |
| Pre vs. During | 1.391 | 0.9345 to 1.847 | Yes | **** | <0.0001 | A-B |  |  |
| Pre vs. Post 1 | 1.065 | 0.6084 to 1.521 | Yes | **** | <0.0001 | A-C |  |  |
| Pre vs. Post 2 | -0.4038 | -0.8600 to 0.05247 | No | ns | 0.0894 | A-D |  |  |
| During vs. Post 1 | -0.3260 | -0.7823 to 0.1302 | No | ns | 0.2012 | B-C |  |  |
| During vs. Post 2 | -1.795 | -2.251 to -1.338 | Yes | **** | <0.0001 | B-D |  |  |
| Post 1 vs. Post 2 | -1.468 | -1.925 to -1.012 | Yes | **** | <0.0001 | C-D |  |  |
| Test details | Mean 1 | Mean 2 | Mean Diff. | SE of diff. | n1 | n2 | q | DF |
| Pre vs. During | -0.0003645 | -1.391 | 1.391 | 0.1537 | 4 | 4 | 12.80 | 12 |
| Pre vs. Post 1 | -0.0003645 | -1.065 | 1.065 | 0.1537 | 4 | 4 | 9.798 | 12 |
| Pre vs. Post 2 | -0.0003645 | 0.4034 | -0.4038 | 0.1537 | 4 | 4 | 3.716 | 12 |
| During vs. Post 1 | -1.391 | -1.065 | -0.3260 | 0.1537 | 4 | 4 | 3.000 | 12 |
| During vs. Post 2 | -1.391 | 0.4034 | -1.795 | 0.1537 | 4 | 4 | 16.51 | 12 |
| Post 1 vs. Post 2 | -1.065 | 0.4034 | -1.468 | 0.1537 | 4 | 4 | 13.51 | 12 |

**Figure 2F**

| Number of families | 1 |  |  |  |  |  |  |  |
| --- | --- | --- | --- | --- | --- | --- | --- | --- |
| Number of comparisons per family | 6 |  |  |  |  |  |  |  |
| Alpha | 0.05 |  |  |  |  |  |  |  |
| Tukey's multiple comparisons test | Mean Diff. | 95.00% CI of diff. | Significant? | Summary | Adjusted P Value |  |  |  |
| Pre vs. During | 21.00 | 10.08 to 31.91 | Yes | *** | 0.0005 | A-B |  |  |
| Pre vs. Post 1 | 21.00 | 10.08 to 31.91 | Yes | *** | 0.0005 | A-C |  |  |
| Pre vs. Post 2 | -27.07 | -37.98 to -16.15 | Yes | **** | <0.0001 | A-D |  |  |
| During vs. Post 1 | 0.000 | -10.92 to 10.92 | No | ns | >0.9999 | B-C |  |  |
| During vs. Post 2 | -48.06 | -58.98 to -37.15 | Yes | **** | <0.0001 | B-D |  |  |
| Post 1 vs. Post 2 | -48.06 | -58.98 to -37.15 | Yes | **** | <0.0001 | C-D |  |  |
| Test details | Mean 1 | Mean 2 | Mean Diff. | SE of diff. | n1 | n2 | q | DF |
| Pre vs. During | 21.00 | 0.000 | 21.00 | 3.676 | 4 | 4 | 8.077 | 12 |
| Pre vs. Post 1 | 21.00 | 0.000 | 21.00 | 3.676 | 4 | 4 | 8.077 | 12 |
| Pre vs. Post 2 | 21.00 | 48.06 | -27.07 | 3.676 | 4 | 4 | 10.41 | 12 |
| During vs. Post 1 | 0.000 | 0.000 | 0.000 | 3.676 | 4 | 4 | 0.000 | 12 |
| During vs. Post 2 | 0.000 | 48.06 | -48.06 | 3.676 | 4 | 4 | 18.49 | 12 |
| Post 1 vs. Post 2 | 0.000 | 48.06 | -48.06 | 3.676 | 4 | 4 | 18.49 | 12 |

**Figure 3D**

| Compare column means (main column effect) |  |  |  |  |  |  |  |  |
| --- | --- | --- | --- | --- | --- | --- | --- | --- |
| Number of families | 1 |  |  |  |  |  |  |  |
| Number of comparisons per family | 15 |  |  |  |  |  |  |  |
| Alpha | 0.05 |  |  |  |  |  |  |  |
| Tukey's multiple comparisons test | Mean Diff. | 95.00% CI of diff. | Significant? | Summary | Adjusted P Value |  |  |  |
| Sevo vs. Control | 129.9 | 88.42 to 171.3 | Yes | **** | <0.0001 |  |  |  |
| Sevo vs. Swim | 54.69 | 13.23 to 96.14 | Yes | ** | 0.0070 |  |  |  |
| Sevo vs. Spry | 116.1 | 74.62 to 157.5 | Yes | **** | <0.0001 |  |  |  |
| Sevo vs. Attack | 129.7 | 88.23 to 171.1 | Yes | **** | <0.0001 |  |  |  |
| Sevo vs. Restraint | 108.5 | 67.05 to 150.0 | Yes | **** | <0.0001 |  |  |  |
| Control vs. Swim | -75.19 | -116.7 to -33.73 | Yes | *** | 0.0004 |  |  |  |
| Control vs. Spry | -13.80 | -55.26 to 27.66 | No | ns | 0.8812 |  |  |  |
| Control vs. Attack | -0.1875 | -41.65 to 41.27 | No | ns | >0.9999 |  |  |  |
| Control vs. Restraint | -21.37 | -62.83 to 20.09 | No | ns | 0.5667 |  |  |  |
| Swim vs. Spry | 61.39 | 19.93 to 102.8 | Yes | ** | 0.0026 |  |  |  |
| Swim vs. Attack | 75.01 | 33.55 to 116.5 | Yes | *** | 0.0004 |  |  |  |
| Swim vs. Restraint | 53.82 | 12.37 to 95.28 | Yes | ** | 0.0079 |  |  |  |
| Spry vs. Attack | 13.61 | -27.84 to 55.07 | No | ns | 0.8869 |  |  |  |
| Spry vs. Restraint | -7.567 | -49.03 to 33.89 | No | ns | 0.9900 |  |  |  |
| Attack vs. Restraint | -21.18 | -62.64 to 20.28 | No | ns | 0.5753 |  |  |  |
| Test details | Mean 1 | Mean 2 | Mean Diff. | SE of diff. | N1 | N2 | q | DF |
| Sevo vs. Control | 139.6 | 9.750 | 129.9 | 12.76 | 4 | 4 | 14.39 | 15.00 |
| Sevo vs. Swim | 139.6 | 84.94 | 54.69 | 12.76 | 4 | 4 | 6.061 | 15.00 |
| Sevo vs. Spry | 139.6 | 23.55 | 116.1 | 12.76 | 4 | 4 | 12.86 | 15.00 |
| Sevo vs. Attack | 139.6 | 9.937 | 129.7 | 12.76 | 4 | 4 | 14.37 | 15.00 |
| Sevo vs. Restraint | 139.6 | 31.12 | 108.5 | 12.76 | 4 | 4 | 12.03 | 15.00 |
| Control vs. Swim | 9.750 | 84.94 | -75.19 | 12.76 | 4 | 4 | 8.333 | 15.00 |
| Control vs. Spry | 9.750 | 23.55 | -13.80 | 12.76 | 4 | 4 | 1.530 | 15.00 |
| Control vs. Attack | 9.750 | 9.937 | -0.1875 | 12.76 | 4 | 4 | 0.02078 | 15.00 |
| Control vs. Restraint | 9.750 | 31.12 | -21.37 | 12.76 | 4 | 4 | 2.368 | 15.00 |
| Swim vs. Spry | 84.94 | 23.55 | 61.39 | 12.76 | 4 | 4 | 6.804 | 15.00 |
| Swim vs. Attack | 84.94 | 9.937 | 75.01 | 12.76 | 4 | 4 | 8.313 | 15.00 |
| Swim vs. Restraint | 84.94 | 31.12 | 53.82 | 12.76 | 4 | 4 | 5.965 | 15.00 |
| Spry vs. Attack | 23.55 | 9.937 | 13.61 | 12.76 | 4 | 4 | 1.509 | 15.00 |
| Spry vs. Restraint | 23.55 | 31.12 | -7.567 | 12.76 | 4 | 4 | 0.8387 | 15.00 |
| Attack vs. Restraint | 9.937 | 31.12 | -21.18 | 12.76 | 4 | 4 | 2.347 | 15.00 |

**Figure 3E**

| Compare column means (main column effect) |  |  |  |  |  |  |  |  |
| --- | --- | --- | --- | --- | --- | --- | --- | --- |
| Number of families | 1 |  |  |  |  |  |  |  |
| Number of comparisons per family | 15 |  |  |  |  |  |  |  |
| Alpha | 0.05 |  |  |  |  |  |  |  |
| Tukey's multiple comparisons test | Mean Diff. | 95.00% CI of diff. | Significant? | Summary | Adjusted P Value |  |  |  |
| Sevo vs. Control | 129.9 | 88.42 to 171.3 | Yes | **** | <0.0001 |  |  |  |
| Sevo vs. Swim | 54.69 | 13.23 to 96.14 | Yes | ** | 0.0070 |  |  |  |
| Sevo vs. Spry | 116.1 | 74.62 to 157.5 | Yes | **** | <0.0001 |  |  |  |
| Sevo vs. Attack | 129.7 | 88.23 to 171.1 | Yes | **** | <0.0001 |  |  |  |
| Sevo vs. Restraint | 108.5 | 67.05 to 150.0 | Yes | **** | <0.0001 |  |  |  |
| Control vs. Swim | -75.19 | -116.7 to -33.73 | Yes | *** | 0.0004 |  |  |  |
| Control vs. Spry | -13.80 | -55.26 to 27.66 | No | ns | 0.8812 |  |  |  |
| Control vs. Attack | -0.1875 | -41.65 to 41.27 | No | ns | >0.9999 |  |  |  |
| Control vs. Restraint | -21.37 | -62.83 to 20.09 | No | ns | 0.5667 |  |  |  |
| Swim vs. Spry | 61.39 | 19.93 to 102.8 | Yes | ** | 0.0026 |  |  |  |
| Swim vs. Attack | 75.01 | 33.55 to 116.5 | Yes | *** | 0.0004 |  |  |  |
| Swim vs. Restraint | 53.82 | 12.37 to 95.28 | Yes | ** | 0.0079 |  |  |  |
| Spry vs. Attack | 13.61 | -27.84 to 55.07 | No | ns | 0.8869 |  |  |  |
| Spry vs. Restraint | -7.567 | -49.03 to 33.89 | No | ns | 0.9900 |  |  |  |
| Attack vs. Restraint | -21.18 | -62.64 to 20.28 | No | ns | 0.5753 |  |  |  |
| Test details | Mean 1 | Mean 2 | Mean Diff. | SE of diff. | N1 | N2 | q | DF |
| Sevo vs. Control | 139.6 | 9.750 | 129.9 | 12.76 | 4 | 4 | 14.39 | 15.00 |
| Sevo vs. Swim | 139.6 | 84.94 | 54.69 | 12.76 | 4 | 4 | 6.061 | 15.00 |
| Sevo vs. Spry | 139.6 | 23.55 | 116.1 | 12.76 | 4 | 4 | 12.86 | 15.00 |
| Sevo vs. Attack | 139.6 | 9.937 | 129.7 | 12.76 | 4 | 4 | 14.37 | 15.00 |
| Sevo vs. Restraint | 139.6 | 31.12 | 108.5 | 12.76 | 4 | 4 | 12.03 | 15.00 |
| Control vs. Swim | 9.750 | 84.94 | -75.19 | 12.76 | 4 | 4 | 8.333 | 15.00 |
| Control vs. Spry | 9.750 | 23.55 | -13.80 | 12.76 | 4 | 4 | 1.530 | 15.00 |
| Control vs. Attack | 9.750 | 9.937 | -0.1875 | 12.76 | 4 | 4 | 0.02078 | 15.00 |
| Control vs. Restraint | 9.750 | 31.12 | -21.37 | 12.76 | 4 | 4 | 2.368 | 15.00 |
| Swim vs. Spry | 84.94 | 23.55 | 61.39 | 12.76 | 4 | 4 | 6.804 | 15.00 |
| Swim vs. Attack | 84.94 | 9.937 | 75.01 | 12.76 | 4 | 4 | 8.313 | 15.00 |
| Swim vs. Restraint | 84.94 | 31.12 | 53.82 | 12.76 | 4 | 4 | 5.965 | 15.00 |
| Spry vs. Attack | 23.55 | 9.937 | 13.61 | 12.76 | 4 | 4 | 1.509 | 15.00 |
| Spry vs. Restraint | 23.55 | 31.12 | -7.567 | 12.76 | 4 | 4 | 0.8387 | 15.00 |
| Attack vs. Restraint | 9.937 | 31.12 | -21.18 | 12.76 | 4 | 4 | 2.347 | 15.00 |

**Figure 3G**

| Compare column means (main column effect) |  |  |  |  |  |  |  |  |
| --- | --- | --- | --- | --- | --- | --- | --- | --- |
| Number of families | 1 |  |  |  |  |  |  |  |
| Number of comparisons per family | 15 |  |  |  |  |  |  |  |
| Alpha | 0.05 |  |  |  |  |  |  |  |
| Tukey's multiple comparisons test | Mean Diff. | 95.00% CI of diff. | Significant? | Summary | Adjusted P Value |  |  |  |
| Sevo vs. Control | -0.08125 | -0.4776 to 0.3151 | No | ns | 0.9832 |  |  |  |
| Sevo vs. Swim | -0.2188 | -0.6151 to 0.1776 | No | ns | 0.4979 |  |  |  |
| Sevo vs. Spry | -0.5063 | -0.9026 to -0.1099 | Yes | ** | 0.0091 |  |  |  |
| Sevo vs. Attack | -0.4938 | -0.8901 to -0.09741 | Yes | * | 0.0110 |  |  |  |
| Sevo vs. Restraint | -0.5313 | -0.9276 to -0.1349 | Yes | ** | 0.0061 |  |  |  |
| Control vs. Swim | -0.1375 | -0.5338 to 0.2588 | No | ns | 0.8627 |  |  |  |
| Control vs. Spry | -0.4250 | -0.8213 to -0.02866 | Yes | * | 0.0322 |  |  |  |
| Control vs. Attack | -0.4125 | -0.8088 to -0.01616 | Yes | * | 0.0391 |  |  |  |
| Control vs. Restraint | -0.4500 | -0.8463 to -0.05366 | Yes | * | 0.0219 |  |  |  |
| Swim vs. Spry | -0.2875 | -0.6838 to 0.1088 | No | ns | 0.2320 |  |  |  |
| Swim vs. Attack | -0.2750 | -0.6713 to 0.1213 | No | ns | 0.2708 |  |  |  |
| Swim vs. Restraint | -0.3125 | -0.7088 to 0.08384 | No | ns | 0.1674 |  |  |  |
| Spry vs. Attack | 0.01250 | -0.3838 to 0.4088 | No | ns | >0.9999 |  |  |  |
| Spry vs. Restraint | -0.02500 | -0.4213 to 0.3713 | No | ns | >0.9999 |  |  |  |
| Attack vs. Restraint | -0.03750 | -0.4338 to 0.3588 | No | ns | 0.9995 |  |  |  |
| Test details | Mean 1 | Mean 2 | Mean Diff. | SE of diff. | N1 | N2 | q | DF |
| Sevo vs. Control | 0.2188 | 0.3000 | -0.08125 | 0.1220 | 4 | 4 | 0.9419 | 15.00 |
| Sevo vs. Swim | 0.2188 | 0.4375 | -0.2188 | 0.1220 | 4 | 4 | 2.536 | 15.00 |
| Sevo vs. Spry | 0.2188 | 0.7250 | -0.5063 | 0.1220 | 4 | 4 | 5.869 | 15.00 |
| Sevo vs. Attack | 0.2188 | 0.7125 | -0.4938 | 0.1220 | 4 | 4 | 5.724 | 15.00 |
| Sevo vs. Restraint | 0.2188 | 0.7500 | -0.5313 | 0.1220 | 4 | 4 | 6.159 | 15.00 |
| Control vs. Swim | 0.3000 | 0.4375 | -0.1375 | 0.1220 | 4 | 4 | 1.594 | 15.00 |
| Control vs. Spry | 0.3000 | 0.7250 | -0.4250 | 0.1220 | 4 | 4 | 4.927 | 15.00 |
| Control vs. Attack | 0.3000 | 0.7125 | -0.4125 | 0.1220 | 4 | 4 | 4.782 | 15.00 |
| Control vs. Restraint | 0.3000 | 0.7500 | -0.4500 | 0.1220 | 4 | 4 | 5.217 | 15.00 |
| Swim vs. Spry | 0.4375 | 0.7250 | -0.2875 | 0.1220 | 4 | 4 | 3.333 | 15.00 |
| Swim vs. Attack | 0.4375 | 0.7125 | -0.2750 | 0.1220 | 4 | 4 | 3.188 | 15.00 |
| Swim vs. Restraint | 0.4375 | 0.7500 | -0.3125 | 0.1220 | 4 | 4 | 3.623 | 15.00 |
| Spry vs. Attack | 0.7250 | 0.7125 | 0.01250 | 0.1220 | 4 | 4 | 0.1449 | 15.00 |
| Spry vs. Restraint | 0.7250 | 0.7500 | -0.02500 | 0.1220 | 4 | 4 | 0.2898 | 15.00 |
| Attack vs. Restraint | 0.7125 | 0.7500 | -0.03750 | 0.1220 | 4 | 4 | 0.4347 | 15.00 |
